# The mechanisms of dynamin-actin interaction

**DOI:** 10.1101/586461

**Authors:** Ruihui Zhang, Nathalie Gerassimov, Donghoon M. Lee, John R. Jimah, Sangjoon Kim, Delgermaa Luvsanjav, Jonathan Winkelman, Marcel Mettlen, Michael E. Abrams, Raghav Kalia, Peter Keene, Pratima Pandey, Benjamin Ravaux, Ji Hoon Kim, Jonathon A. Ditlev, Guofeng Zhang, Michael K. Rosen, Adam Frost, Neal M. Alto, Margaret Gardel, Sandra L. Schmid, Jenny E. Hinshaw, Elizabeth H. Chen

**Author notes:** These authors contributed equally (Ruihui Zhang, Nathalie Gerassimov, Donghoon M. Lee, John R. Jimah). Correspondence and requests for materials should be addressed to E.H.C.

## Abstract

Cell-cell fusion is an indispensable process in the conception, development and physiology of multicellular organisms. Here we demonstrate a direct and noncanonical role for dynamin, best known as a fission GTPase in endocytosis, in cell-cell fusion. Our genetic and cell biological analyses show that dynamin colocalizes within the F-actin-enriched podosome-like structures at the fusogenic synapse, which is required for generating invasive membrane protrusions and myoblast fusion in vivo, in an endocytosis-independent manner. Biochemical, negative stain EM and cryo-electron tomography (cryo-ET) analyses revealed that dynamin forms helices that directly bundles actin filaments by capturing multiple actin filaments at their outer rim via interactions with dynamin’s proline-rich domain. GTP hydrolysis by dynamin triggers disassembly of the dynamin helix, exposes the sides of the actin filaments, promotes dynamic Arp2/3-mediated branched actin polymerization, and generates a mechanically stiff actin network. Thus, dynamin functions as a unique actin-bundling protein that enhances mechanical force generation by the F-actin network in a GTPase-dependent manner. Our findings have universal implications for understanding dynamin-actin interactions in various cellular processes beyond cell-cell fusion.

---

Cell-cell fusion is essential for the conception, development, regeneration and physiology of multicellular organisms^1-3^. A common theme underlying cell-cell fusion events ranging from insects to mammals is the presence of actin-propelled invasive protrusions at the site of fusion, known as the fusogenic synapse^4-8^. Such invasive protrusions were first identified as part of an F-actin-enriched podosome-like structure (PLS) in *Drosophila* myoblast fusion^7^. The PLS is generated by an “attacking” fusion partner, the fusion-competent myoblast (FCM), which drills multiple invasive protrusions into the “receiving” fusion partner, the muscle founder cell, to promote cell membrane juxtaposition and fusion pore formation^7,9,10^. Although each protrusion has a diameter similar to that of a filopodium, these protrusions are mechanically stiffer and are capable of triggering mechanosensitive responses in the receiving cell^11^. Mechanoresponsive accumulations of myosin II and spectrin in the receiving cell, in turn, increase the cortical tension of the receiving cell, thus generating resisting forces against the invasive protrusions and bringing the two cell membranes to close proximity to promote cell fusion^11,12^. The invasive protrusions from the attacking cells are propelled by Arp2/3-mediated branched actin polymerization, which generates short actin filaments more suitable for mechanical work than long and linear filaments^9^. However, how the branched actin filaments are organized to generate mechanically stiff membrane protrusions during fusion is unknown.

Dynamin (Dyn) is a large GTPase best known for its role in endocytosis. Dynamin catalyzes membrane fission by forming rings/helices around necks of a budding endocytic vesicles^13-15^. The dynamin protein consists of an amino-terminal G domain that catalyzes GTP hydrolysis, a three helix bundle-signaling elements (BSEs), a stalk critical for oligomerization, a pleckstrin homology (PH) domain that interacts with phosphatidyl-inositol-4,5-biphosphate (PIP2) in the plasma membrane, and a carboxy-terminal proline-rich domain (PRD) for SH3 domain interactions that support proper dynamin localization^13-15^. Crystal structures of dynamin revealed a hairpin-like folding of each monomeric protein mediated by the stalk regions and the BSEs^16,17^. When assembled into rings or helices, the PH domain at the vertex of the hairpin is positioned inside the dynamin ring, directly interacting with the enclosed membrane tubule that is wrapped inside, whereas the G domain is localized at the outer rim of the dynamin helix. The stalk regions form the backbone of the helix in between the PH and the G domains. The unstructured PRD has not been observed in high-resolution structure, and is predicted to be neighboring the G domain and extend outward beyond the G domain^13,18^.

In addition to endocytosis, dynamin regulates the formation and/or function of F-actin-enriched cellular structures, such as podosomes^19,20^, invadopodia^21^, filopodia^22^, lamellipodia^23^, actin comet tails^24,25^, phagocytic cups^26^, and stress fibers^27,28^. While mechanisms underlying dynamin-mediated endocytosis have been extensively studied, how dynamin interacts with actin remains a widely-documented but poorly understood phenomenon. Studies suggest that dynamin bundles actin either directly^29,30^ or indirectly in a cortactin-dependent manner^22,31,32^. The binding of actin filaments directly by dynamin was proposed to antagonize gelsolin-mediated filament capping thus promoting actin polymerization^29^, whereas the cortactin-mediated dynamin-actin interaction was thought to stabilize F-actin filaments^22,31,32^. In either model, the precise mode and functions of dynamin-actin interactions remain unclear.

## Results

### Dynamin is required for myoblast fusion in vivo and colocalizes with the F-actin foci at the fusogenic synapse

The presence of a podosome-like structure at the fusogenic synapse^7^ prompted us to examine whether dynamin, which was implicated in podosome formation^19^, plays a role in *Drosophila* myoblast fusion. The *Drosophila* genome encodes one dynamin, named *shibire (shi)*, which shares 67% protein sequence identity with human dynamins. The availability of temperature-sensitive (ts) alleles of *shi* allowed us to eliminate both maternal and zygotic Shi function in a temporally controlled manner. We incubated the *shi^ts^* embryos at permissive temperature (18°C) for 14.5 hours just prior to the start of muscle development, before shifting to restrictive temperature (34°C) for 4 hours and analyzing the resulting muscle phenotype. Both *shi^ts1^* and *shi^ts2^*, which contain single point mutations in the G domain, exhibited a severe myoblast fusion defect at 34°C characterized by the presence of many unfused, mononucleated myoblasts (Fig. 1A, B; Fig. S1A, B, C). These myoblasts expressed tropomyosin and muscle myosin heavy chain (MHC), indicating that they had properly differentiated (Fig. 1A; Fig. S1A, B). Expressing wild-type Shi-GFP in *shi^ts2^* mutant embryos with an all muscle cells *twi-GAL4* driver or specifically in FCMs with an *sns-GAL4* driver significantly rescued myoblast fusion at 34°C (Fig. 1B; Fig. S1D). In contrast, expressing Shi-GFP specifically in founder cells with the *rP298-GAL4* driver did not rescue the fusion defect. These data establish that the fusion defect was caused specifically by the loss of Shi, especially in the FCMs.

**Fig. 1.**
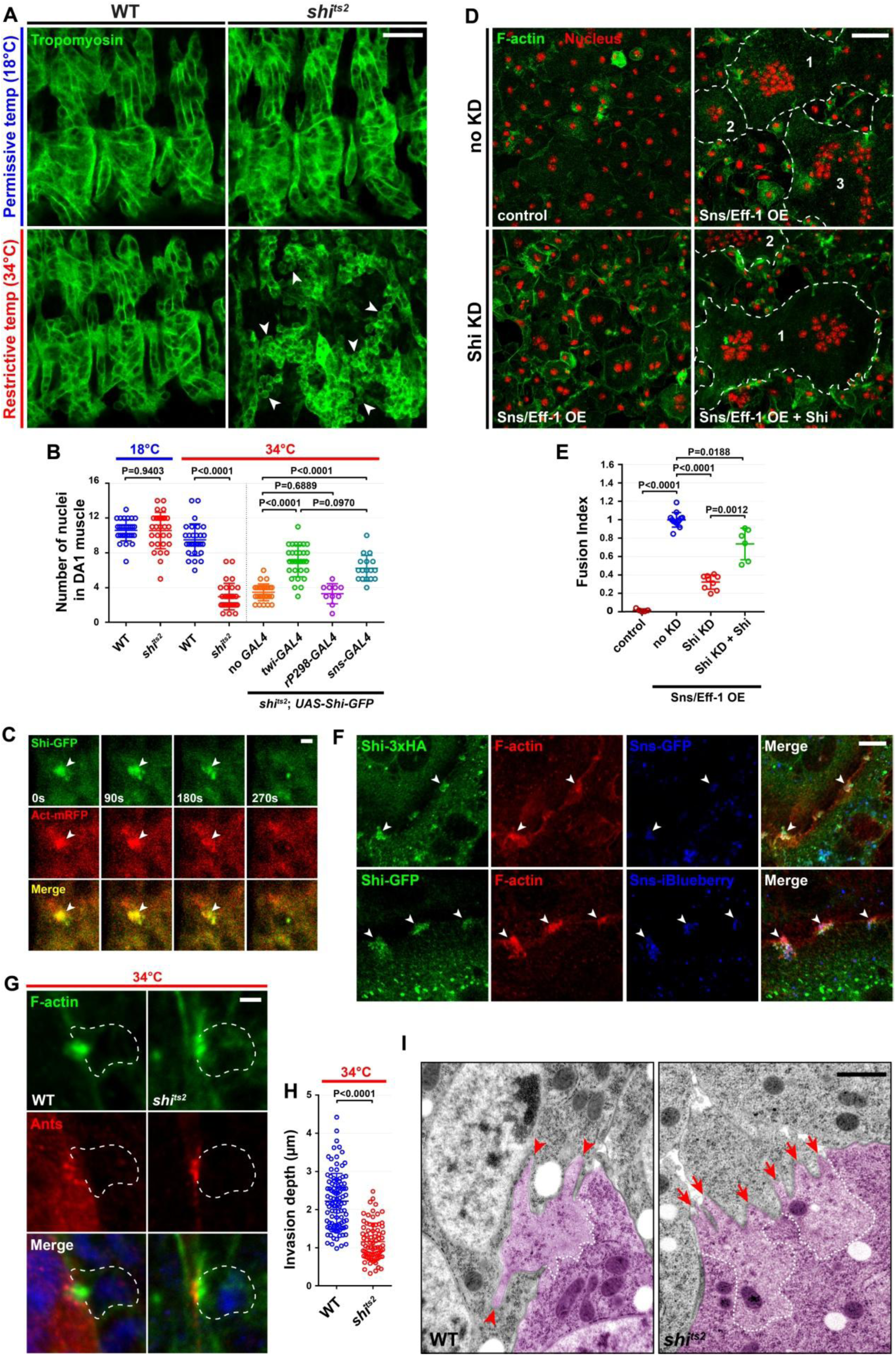
Shibire promotes myoblast fusion by upregulating PLS invasion at the fusogenic synapse. (A) Myoblast fusion is defective in *shi* mutant embryo. The somatic muscles in stage 15 wild-type (WT) and *shi^ts2^* mutant embryos were labeled by anti-Tropomyosin antibody. Note the presence of unfused mononucleated myoblasts in *shi^ts2^* mutant embryo at the restrictive temperature (arrowheads), compared to the normal musculature of *shi^ts2^*mutant embryos at permissive temperature and of WT embryos at both temperatures. Scale bar: 20 µm. (B) Quantification of the fusion index in *Drosophila* embryos. The numbers of Eve-positive nuclei in the dorsal acute muscle 1 (DA1) of stage 15 embryos were counted for each genotype in (A) and Fig S1D. The numbers of DA1 muscles analyzed for each genotype: n = 30, 30, 30, 30, 30, 30, 10 and 17 (left to right). The wide horizontal bars indicate the mean values. Significance was determined by the two-tailed Student’s t-test. Note that myoblast fusion defect in *shi^ts2^* mutant embryos was rescued by expression of Shi-GFP in all muscle cells (*twi-GAL4*) or FCMs (*sns-GAL4*), but not in founder cells (*rP298-GAL4*). (C) Shi colocalizes with F-actin foci at fusogenic synapses in *Drosophila* embryos. Time-lapse stills of a stage 14 embryo expressing Shi-GFP and Actin (Act)-mRFP (see Movie 1). Note that Shi-GFP was enriched in the actin focus (arrowheads) and dissolved together with the actin focus upon myoblast fusion. Scale bar: 2 µm. (D) Shi promotes the fusion of S2R+ cells co-expressing Sns and Eff-1. Sns/Eff-1-expresssing S2R+ cells fused to form multinucleated (≥ 3 nuclei) syncytia (outline and numbered). Cells were labeled with phalloidin (F-actin; green) and Hoechst (nuclei; red). Shi knockdown (KD) (dsRNA targeting Shi 5’UTR) inhibited cell fusion (bi-nucleated cells due to incomplete cytokinesis), which was rescued by expressing full-length Shi lacking the 5’UTR. Scale bar: 50 µm. (E) Quantification of the fusion index in S2R+ cells. The fusion index was determined as the ratio between the number of nuclei in multinucleated syncytia (≥3 nuclei) vs. in total transfected cells, normalized to (the average of) S2R+ cells overexpressing (OE) Sns and Eff-1 within the same set of experiments. Fusion indexes were calculated for experiments in (D). The numbers of independent experiments performed: n = 5, 13, 9, and 6 (left to right). The wide horizontal bars indicate the mean values. Significance was determined by the two-tailed Student’s t-test. (F) Shi colocalizes with F-actin foci at fusogenic synapses in S2R+ cells. Cell expressing Shi-3xHA (green), Sns-GFP (blue) and Eff-1 were labeled with phalloidin (F-actin; red) and anti-HA antibody (top panels). Cells expressing Shi-GFP (green), Sns-iBlueberry (blue) and Eff-1 were labeled with phalloidin (bottom panels). Note that both Shi-3xHA and Shi-GFP colocalized with F-actin at fusogenic synapses indicated by Sns enrichment (arrowheads). Scale bar: 10 µm. (G) The F-actin foci at fusogenic synapses are less invasive in *shi^ts2^* mutant embryo. Stage 14 wild-type (WT) and *shi^ts2^*mutant embryos stained with phalloidin (F-actin; green), anti-antisocial (Ants; founder cell-specific adaptor protein enriched at the fusogenic synapse; red), and Hoechst 33342(nuclear marker; blue) at restrictive temperature. FCMs are outlined. Note the inward curvature reflective of deep actin foci invasion revealed by Ants in WT, but not *shi^ts2^* mutant embryo, corresponding with the deep and shallow F-actin foci invasion depth, respectively. Scale bar: 2 µm. (H) Quantification of F-actin foci invasion depth in WT and *shi^ts2^* mutant embryos. The invasion depths of 102 WT and 76 *shi^ts2^*mutant F-actin foci were measured at restrictive temperature. The thick horizontal bars indicate the mean values. Significance was determined by the two-tailed Student’s t-test. (I) Electron micrographs of the fusogenic synapse in stage 14 WT and *shi*^ts2^ mutant embryos. FCMs are pseudo-colored in purple. Dashed lines delineate the F-actin-enriched area of the PLS. Note long invasive finger-like protrusions in WT (arrowheads) and short and spread out protrusions in the *shi*^ts2^ mutant embryo (arrows). Scale bar: 1 µm.

Consistent with Shi’s role in FCMs, Shi colocalized within the FCM-specific F-actin foci at the fusogenic synapse, visualized by both anti-Shi antibody and muscle-specific expression of Shi-GFP (Fig. 1C; Fig. S2A). Live imaging of embryos coexpressing Shi-GFP and actin-mRFP in muscle cells revealed that Shi colocalized with the F-actin foci during their entire life span and dissolved with the F-actin foci when fusion was completed (Fig. 1C; Movie 1). Recruitment of Shi to the fusogenic synapse dependent on the FCM-specific cell adhesion molecule Sns, but not on the actin nucleation-promoting factors (NPFs) for the Arp2/3 complex, including WASP, the WASP-interacting protein (WIP) or Scar (Fig. S2B-E).

A general role for Shi in cell-cell fusion was supported by experiments in *Drosophila* S2R+ cells that were induced to fuse by co-expressing Sns and a *C. elegans* fusogen Eff-1^8^. Knocking down (KD) Shi using dsRNAs against the 5’ untranslated region (UTR) led to a fusion defect in S2R+ cell fusion, which was significantly rescued by wild-type Shi expression (Fig. 1D, E). As in *Drosophila* embryos, Shi colocalized with F-actin foci at the fusogenic synapse in S2R+ cells (Fig. 1F), consistent with a specific function of Shi at the fusogenic synapse. Although a role for dynamin 2 (Dyn2) in the fusion of cultured mouse myoblasts and osteoblasts has been reported in previous studies, the underlying mechanisms have been attributed to either dynamin-mediated endocytosis or fusion pore expansion^5,33^. The nature of any direct, physical associations between dynamin and the actin-enriched structure at the fusogenic synapse have not been investigated.

### Dynamin is required for generating invasive protrusions at the fusogenic synapse independent of endocytosis

In wild-type embryos, founder cells are invaded by the FCM-specific PLS with an average depth of invasion of 2.2 ± 0.7 μm (n=102) (Fig. 1G, left panels; Fig. 1H). However, the PLS in *shi^ts2^* mutant were broadened and had a decreased invasive depth of 1.2 ± 0.5 μm (n=76) at the restrictive temperature (Fig. 1G, right panels; Fig. 1H). Consistent with the observations using confocal microscopy, ultrastructural studies using transmission electron microscopy (TEM) revealed stubby protrusions originating from a broad base of actin enrichment in *shi^ts2^* mutant embryos, as compared to the long finger-like protrusions projecting from a dense actin-enriched area in wild-type embryos (Fig. 1I). These data suggest that Shi is involved in promoting the stiff, elongated finger-like protrusions in wild-type embryos.

If the stubby membrane protrusions in *shi^ts2^* mutant were caused by a defect in endocytosis, one would expect an accumulation of endocytic pits along the cell membranes at the fusogenic synapse at restrictive temperature, as observed at the neuromuscular junction in the *shi^ts1^* mutant^34^. However, no endocytic pits were observed at the fusogenic synapse under the restrictive temperature (Fig. 1I). In addition, neither clathrin light chain nor the *α* subunit of the AP-2 clathrin adaptor complex was enriched at the fusogenic synapse in wild type embryos (Fig. S2F). Thus, clathrin-mediated endocytosis appears not to be associated with the enrichment of Shi at the fusogenic synapse.

### Dynamin bundles actin filaments by forming regularly spaced ring/helices

The colocalization of Shi with the F-actin foci and its role in promoting invasive protrusions prompted us to investigate potential interactions between Shi and actin. Low speed F-actin co-sedimentation assays revealed that purified Shi bundled rabbit skeletal muscle actin in a concentration-dependent manner under low ionic strength solution conditions (50 mM KCl). which is known to allow the formation of higher order structures of dynamin, such as rings and helices^35,36^ (Fig. 2A; Fig. S3A). In addition, high speed co-sedimentation assays showed that Shi bound F-actin (Fig. S3B), and the apparent K_d_ of Shi-actin interaction was 0.19 ± 0.03 μM (Fig. S3C, D), similar to that of human Dyn1-actin interaction (∼ 0.4 μM)^29^.

**Fig. 2.**
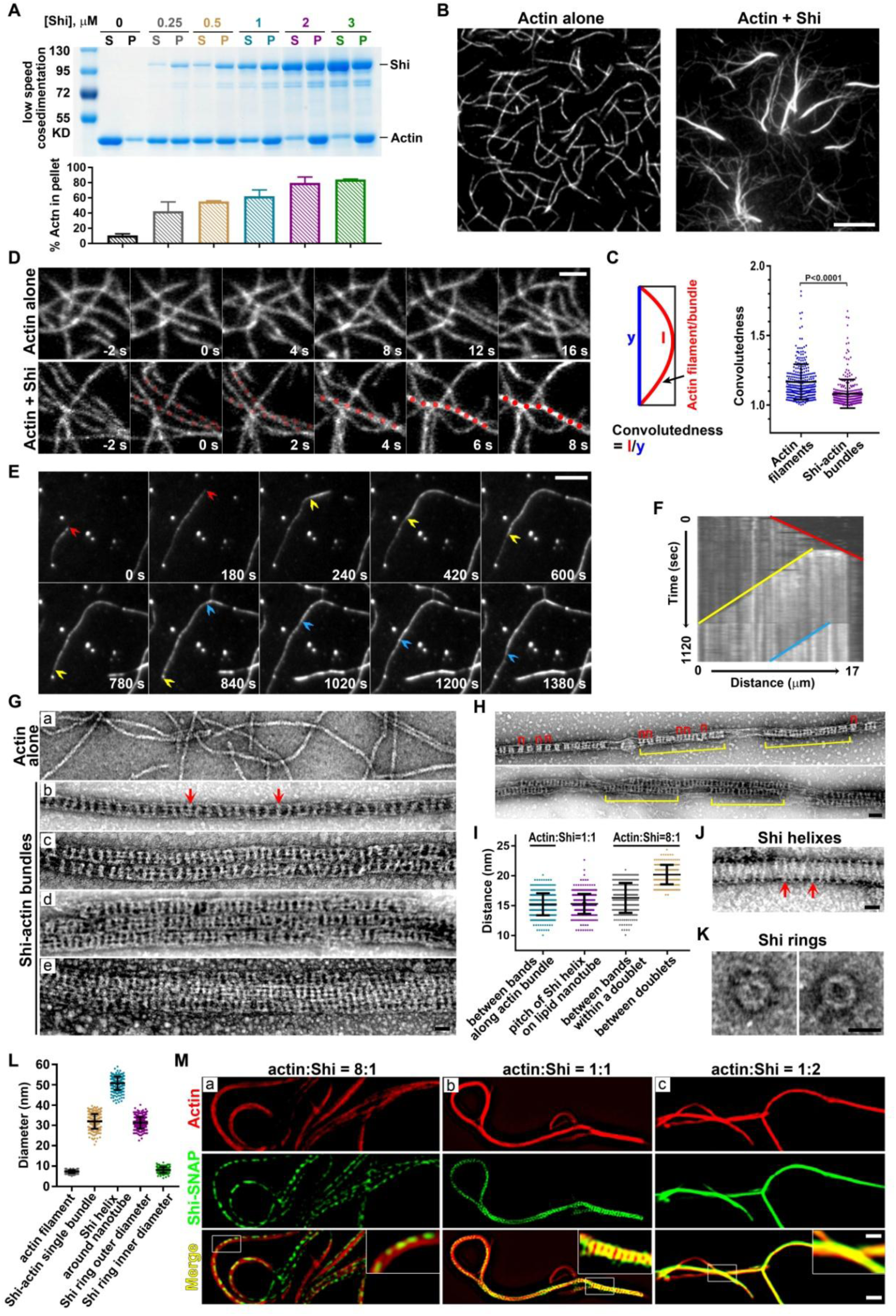
Shibire bundles actin filaments. (A) Shi bundles actin filaments in a dose-dependent manner. In the low speed F-actin co-sedimentation assays, 3 μM actin was incubated with increasing concentrations of Shi, and supernatants (S) and pellets (P) were monitored by SDS-PAGE after low speed centrifugation at 13,600*g*. The percentage of actin (mean±SD, n=3) in the pellet vs. total was quantified. (B) TIRF images showing actin filaments (left panel) and Shi-mediated actin bundles (right panel). The TIRF assay was performed in NEM-myosin-coated flow chambers. Scale bar, 5 μm. (C) Schematic diagram of the measurement of convolutedness (left), which is defined as the ratio of traced filament length to the length of the longest side of a bounding rectangle encompassing the same filament. Plot of the convolutedness of actin filaments and Shi-mediated actin bundles in (B) (right). Data represent measurements (mean±SD) of 328 actin filaments and 304 Shi-mediated actin bundles. Significance was determined by two-tailed Student’s *t*-test. (D) Time-lapse stills of TIRF images showing Shi-mediated actin bundling (see Movie 2). The TIRF assay was performed on supported lipid bilayers. The bundled actin filaments are indicated by red dots. Scale bar, 2 μm. (E) Time-lapse stills of TIRF images showing the process of actin bundling during filament growth (see Movie 3). The TIRF assay was performed in mPeg-Silane-coated flow chambers. The fast-growing ends of three actin filaments are indicated by red, yellow and blue arrowheads, respectively. Scale bar, 5 μm. (F) Kymograph of the actin bundle formation shown in (E). Three colored lines indicate the tracks of fast-growing ends of the first (red), second (yellow) and third (blue) filaments. (G) Transmission electron micrographs of negatively stained actin filaments (a) and Shi-mediated actin bundles (b-e) ([actin]:[Shi] = 1:1). (b) A single Shi-actin bundle. Shi rings appeared as regularly spaced, light-colored bands along the actin bundle (arrows). (c) A double bundle. Note the largely un-synchronized Shi rings of the two closely aligned bundles. (d) A triple bundle with unaligned Shi rings. (e) A multi-bundle with mostly aligned Shi rings across the bundles. Scale bar, 30 nm. (H) Transmission electron micrographs of negatively stained single and double Shi-actin bundles ([actin]:[Shi] = 8:1). Note that Shi rings are arranged in doublets indicated by red brackets, and the presence of segments of “naked” actin filaments not associated with stretches of Shi rings (yellow brackets). Scale bar, 50 nm. (I) Dot plot of the distance between Shi rings ([actin]:[Shi] = 1:1) along the actin bundle and the pitch of Shi helices on lipid nanotube shown in (J), and the distance between Shi rings within and between the Shi doublets ([actin]:[Shi] = 8:1). Values in the bar graph represent mean±SD, *n* = 401, 331, 315, and 189, respectively. (J) Transmission electron micrographs of negatively stained lipid nanotube and Shi. Arrows indicate Shi helices wrapping around the nanotube. Scale bar, 30 nm. (K) Transmission electron micrographs of negatively stained Shi rings. Scale bar, 30 nm. (L) Dot plot of the average diameters of actin filament, Shi-actin bundle, Shi helices around nanotube, and Shi rings. Values in the bar graph represent mean±SD, n = 219, 259, 252, 282 and 198, respectively. (M) SIM images of actin bundles formed at three different [actin]/[Shi] ratios indicated. Images were collected after a 2-hr incubation of actin labeled with Alexa568-conjugated phalloidin (red) and Shi-SNAP-Surface 488 (green). Boxed areas are enlarged in the insets. Scale bars, 1 μm (main panels) and 0.4 μm (insets).

Using total internal reflection fluorescence (TIRF) microscopy, we demonstrated that actin filaments converge together to form thick actin bundles in the presence of Shi in all three conditions tested: on coverslips coated with N-ethylmaleimide (NEM)-modified myosin fragments (Fig. 2B), on supported lipid bilayers (Fig. 2D; Movie 2), and on coverslips coated with mPEG-Silane (Fig. 2E; Movie 3). The thick actin bundles were significantly straighter compared to single actin filaments, indicating that the bundles were mechanically stiffer (Fig. 2C). When single filaments were imaged in a floe chamber, we observed that once an initial filament was generated, a second filament appeared to grow alongside it to form a bi-filament “mini-bundle”, which was then joined by a third filament growing alongside the bi-filament bundle (Fig. 2E, F; Movie 3).

Negative stain EM revealed that when mixed at equal concentrations (1 μM) Shi formed regularly spaced, lighter staining “bands” along the actin bundles (Fig. 2Ga, b). The average distance between the Shi bands was 15.1 ± 1.8 nm (n= 401), comparable to the average pitch of Shi helices around lipid tubules (15.5 ± 1.6 nm; n= 331) (Fig. 2I, J). We refer to the Shi “bands” as Shi rings, since at this resolution it was difficult to determine whether Shi formed individual rings or helices. Interestingly, at low Shi:actin (0.125 μM Shi:1 μM actin), Shi rings appeared to be arranged in “doublets” along the actin bundle and there were stretches of the actin bundle without Shi rings (Fig. 2H). The average distance between the two Shi rings within a doublet was 16.3 ± 2.5 nm (n= 315), and that between two doublets was 20.2 ± 1.6 nm (n= 189) (Fig. 2I), with the sum of the two distances (∼ 36.5 nm) largely coinciding with a half helical pitch of actin (36 nm). Such characteristic spacing of Shi rings at a low Shi concentration suggests that Shi may have high affinity interactions with specific sites on the actin filaments. In addition to single actin bundles, we also observed “super-bundles” in which two, three or more actin bundles aligned in parallel, each of which could be distinguished by the slightly misaligned Shi rings (Fig. 2Gc, d). Occasionally, the Shi rings appeared more aligned across multiple actin bundles (Fig. 2Ge).

Interestingly, the average outer diameter of an actin bundle was 32.0 ± 3.6 nm (n= 259), which is similar to that of Shi rings alone (31.3 ± 2.9 nm; n= 282) (Fig. 2G, K, L), indicating that bundling actin did not significantly change the size of the Shi ring. This is in contrast to the increased diameter of Shi helices surrounding membrane nanotubes (50.7 ± 3.3 nm; n=252) (Fig. 2J, L). Like *Drosophila* Shi, human Dyn1 and 2 also formed regularly spaced rings that directly bundled actin shown by negative stain EM analysis of negative stained samples (Fig. S4Aa-c and e-f). The average distances between the hDyn1 and hDyn2 rings along actin bundle were 15.6 ± 1.4 nm (n=307) and 15.4 ± 1.4 nm (n=245), respectively (Fig. S4A, B), similar to the average pitch of hDyn1 helices around nanotubes, 15.4 ± 1.7 nm (n=252) (Fig. S4B, E). Moreover, the average outer diameter of the hDyn1 rings along actin bundles (44.7 ± 3.0 nm; n= 220) (Fig. S4A, C) was significantly larger than that of *Drosophila* Shi (32.0 ± 3.6 nm; n= 259) (Fig. 2G, L), but comparable to the outer diameter of hDyn1 rings alone (40.2 ± 4.3 nm; n= 132), and smaller than that of hDyn1 around nanotubes (51.3 ± 1.8 nm; n=226) and constricted liposomes (∼50 nm)^37^ (Fig. S4C, D, E). Thus, while there are similarities in the spacing between dynamin rings on lipid substrates and those associated with F-actin filaments, the differences in ring diameter suggest structural differences in the manner by which dynamin rings interact with actin filaments vs. membrane tubules.

To visualize Shi along actin bundles using light microscopy, we generated C-terminally SNAP-tagged Shi, which showed normal activity in assembling around nanotubes and bundling actin (Fig. S3E). Super resolution microscopy revealed Shi-SNAP along the actin bundles (Fig. 2M). At sub-stoichiometric concentrations (0.125 μM Shi-SNAP:1 μM actin), Shi-SNAP appeared in thin and discontinuous stretches along actin bundles (Fig. 2Ma), consistent with the EM observation in Fig. 2H. At stoichiometric concentrations (1 μM each), Shi-SNAP was more densely distributed along the super-bundle and formed periodically repeated “bands” across the entire width of the bundle (Fig. 2Mb), likely corresponding to the aligned trans-interacting Shi-SNAP stretches observed by EM (Fig. 2Ge). At 2 μM, Shi-SNAP uniformly covered the entire super-bundle (Fig. 2Mc).

### GTP hydrolysis of dynamin disassembles the dynamin ring and loosens the actin bundles

GTP addition significantly decreased the amount of Shi-SNAP associated with the actin super-bundles, presumably by disassembling the Shi rings, leaving sparsely localized Shi-SNAP punctae (Fig. 3A). Under EM, the actin filaments no longer associated with Shi rings and were loosely scattered by the addition of GTP (Fig. 3B). Consistent with a role for GTP hydrolysis in disassembling Shi ring and actin bundles, point mutations in the Shi G domain that disable GTP hydrolysis blocked GTP-induced Shi ring disassembly and actin bundle dissociation. Specifically, Shi-Q35E (corresponding to human Dyn1-Q40E) (Fig. 3B), Shi-G141S (*Drosophila* ts2 mutant; Dyn1-G146S) (Fig. 3C) and Shi-K137A (Dyn1 K142A) (Fig. 3B) all bundled actin filaments as wild-type Shi in the absence of GTP, but they either completely or partially inhibited ring disassembly and actin bundle dissociation upon GTP addition (at the restrictive temperature for Shi-G141S).

**Fig. 3.**
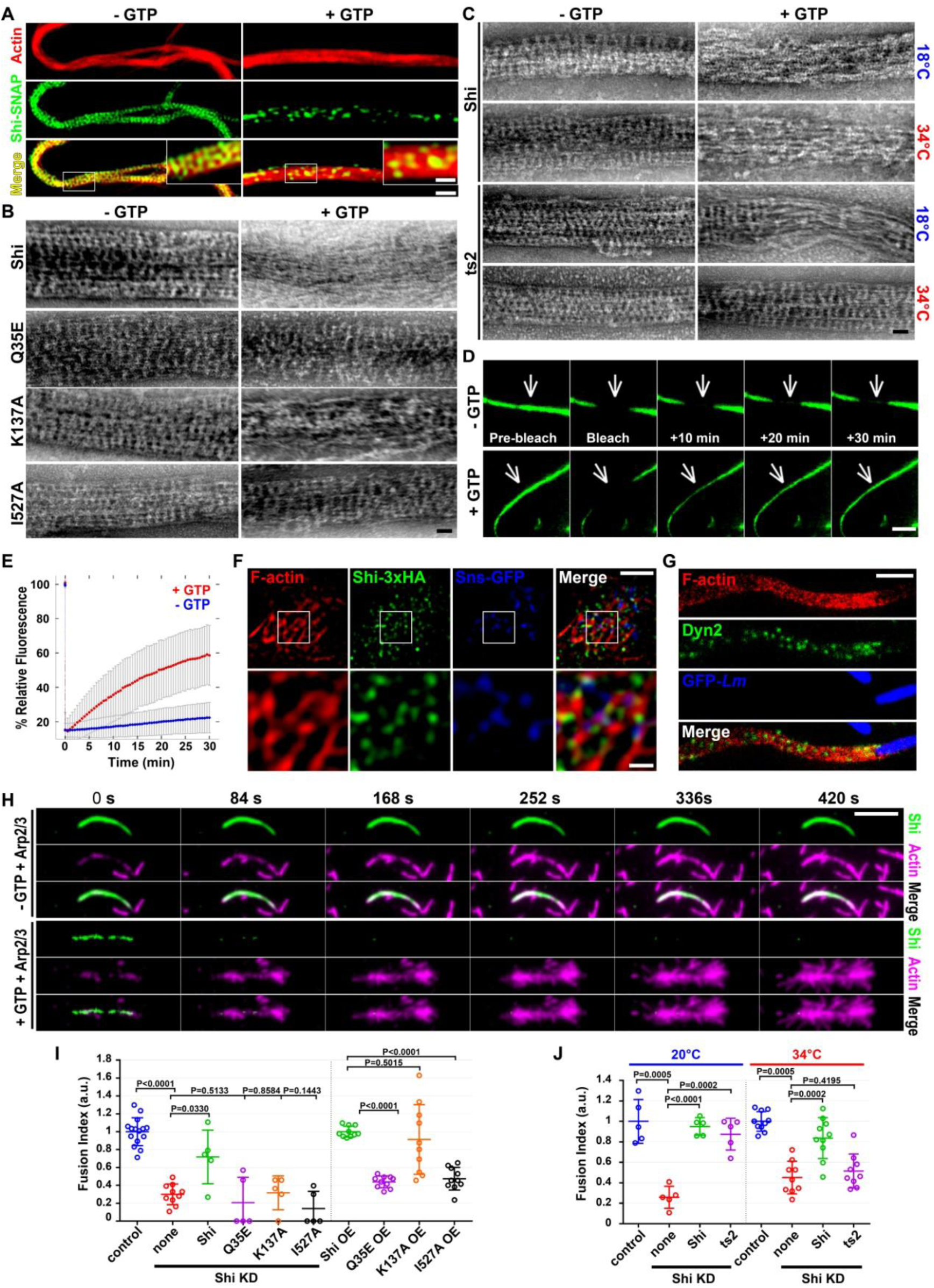
GTP hydrolysis disassembles Shibire rings to allow branched actin polymerization and cell-cell fusion. (A) SIM images of Shi-actin bundles with or without GTP. Images were collected after a 2-hr incubation of actin labeled with Alexa568-conjugated phalloidin (red) and Shi-SNAP-Surface 488 (green) in the presence or absence of GTP. Boxed areas are enlarged in the insets. Note the densely distributed Shi “band” across the actin bundle in the absence of GTP, and the punctate spots of Shi in the presence of GTP. Scale bars, 1 μm (main panels) and 0.4 μm (insets). (B-C) Transmission electron micrographs of negatively stained actin bundles in the absence or presence of GTP. (B) Actin bundles mediated by Shi and the indicated Shi mutants. Wild-type Shi rings were disassembled upon GTP addition, leading to loosened actin filaments (top panel). Q35E and I527A rings failed to disassemble, and K137A rings partially disassembled, upon GTP addition. (C) The ts2 mutant rings disassembled normally at permissive temperature (18°C), but failed to disassemble at the restrictive temperature (34°C), in the presence of GTP. Scale bars, 30 nm. (D) FRAP of Shi-SNAP-Surface 488 on actin bundles in the absence (top panels) or presence (bottom panels) of GTP (see Movie 4). A segment (indicated by arrow) of a Shi-actin bundle was photobleached and the fluorescence recovery was monitored live. Stills of the actin bundle before and after photobleaching are shown. Scale bar, 3 μm. (E) Recovery kinetics of the Shi-SNAP fluorescence after photobleaching in the presence or absence of GTP (n=13 or 9 actin bundles, respectively; grey bars represent SD). (F) Shi localization on actin bundles at the fusogenic synapse revealed by structured illumination microscopy (SIM). S2R+ cells co-expressing Sns-GFP (blue), Eff-1 and Shi-3xHA were labeled with phalloidin (F-actin; red) and anti-HA (green). Boxed areas are enlarged in the insets. Note the Shi punctae and bands along the actin bundles. Scale bars, 2 μm (main panels) or 500 nm (insets). (G) Human Dyn2 localization on actin comet tails in *Listeria-*infected HeLa cells revealed by stimulated emission depletion microscopy (STED). Note the Dyn2 punctae and bands (green) along the actin comet tails labeled with phalloidin (red). *Listeria*-GFP is in blue. Scale bar: 2 µm. (H) Time-lapse stills of TIRF images showing Arp2/3-mediated branched actin polymerization on Shi-actin bundles in the absence or presence of GTP. Shi-actin bundles were generated by incubating unlabeled actin with Shi-SNAP-Surface 488 (green) (see Movie 5). Arp2/3, VCA and rhodamine-actin (magenta) were added to the Shi-actin bundles to start branched actin polymerization without (top panels) or with (bottom panels) 1 mM GTP. Note the disassembly of Shi from the actin bundles upon GTP addition (unlabeled), and the numerous branched actin filaments emanating from the same actin bundles (bottom panels). The increase in the actin bundle intensity in the absence of GTP is likely due to additional bundling (top panels). Scale bar, 5 μm. (I) Quantification of the fusion index in S2R+ cells expressing Shi and various mutants or following dsRNA mediated knockdown (KD) of Shi or when overexpressed (OE), as indicated. Note that in Sns-Eff-1-expressing Shi KD cells, only wild-type Shi, but not the Q35E, K137A and I527A mutants, rescued the cell fusion defect. In Sns-Eff-1-expressing cell, Q35E and I527A OE caused strong dominant negative effect in inhibiting cell fusion, whereas K137A OE did not cause a significant dominant negative effect. All Shi and mutant constructs were tagged with 3xHA. The numbers of independent experiments performed: n = 15, 10, 5, 5, 5, 5, 10, 10, 10 and 10 (left to right). The wide horizontal bars indicate the mean values. Significance was determined by the two-tailed Student’s t-test. (J) Quantification of the fusion index of Shi KD S2R+ cells expressing Shi and ts2 mutant. Note that at the restrictive temperature (34°C), ts2 failed to rescue the fusion defect caused by Shi KD in Sns/Eff-1-expressing S2R+ cells. The numbers of independent experiments performed: n = 5, 5, 5, 5, 10, 9, 10 and 9 (left to right). The wide horizontal bars indicate the mean values. Significance was determined by the two-tailed Student’s t-test.

Fluorescence recovery after photobleaching (FRAP) analyses revealed that actin bundling by Shi in the presence of GTP is a dynamic process. Thus, while little fluorescence recovery was detected after photobleaching a segment of actin-associated Shi-SNAP in the absence of GTP (7.7% ± 8.9%), there was a significant fluorescence recovery (52.0% ± 16.9%) in the presence of GTP. This difference suggests that Shi equilibrates between ring assembly and disassembly when GTP is present (Fig. 3D, E; Movie 4). Similar to Shi, hDyn1 and 2 rings along actin bundles were also disassembled by GTP addition, resulting in loosened actin bundles (Fig. S4Ad, g). We further note that the GTPase activity of Shi was stimulated ∼35 fold in the presence of actin filaments, compared to Shi’s basal activity, consistent with actin-induced, assembly-stimulated hydrolysis (Fig. S6).

### Dynamin bundles actin filaments in vivo

If Shi is involved in bundling actin filaments at the fusogenic synapse, one would expect Shi to colocalize with the actin bundles. However, due to the presence of an average of ∼0.5 μM GTP in cells^38^, one would expect a punctate Shi localization pattern, similar to that of Shi-SNAP on actin super-bundles in the presence of GTP (Fig. 3A). Indeed, super resolution microscopy (SIM and STED) revealed Shi-GFP and Shi-HA punctae decorating the actin bundles at the fusogenic synapse in Sns-Eff-1-expressing S2R+ cells (Fig. 3F; Fig. S5A). To ask whether dynamin exhibits a similar pattern of localization on other actin-enriched structures, such as actin comet tails induced by the bacteria *Listeria monocytogenes*, we examined hDyn2 on comet tails by super resolution microscopy. Indeed, we observed similar punctate hDyn2 localization on actin comet tails in *Listeria-*infected Hela cells (Fig. 3G; Fig. S5B). These data suggest that dynamin is involved in bundling actin filaments in vivo and that GTP hydrolysis allows dynamic assembly and disassembly of dynamin rings along the actin bundles.

### Dynamin ring disassembly is required for Arp2/3-mediated actin polymerization and cell-cell fusion

Given the colocalization of dynamin and the branched actin network at fusogenic synapses and comet tails, we asked how dynamin-actin interaction might regulate Arp2/3-mediated actin polymerization. We reasoned that dynamin rings along the actin bundles could block Arp2/3-actin interaction therefore preventing branched actin polymerization. Indeed, in the absence of GTP, Shi-bound actin bundles failed to initiate Arp2/3-mediated branched actin polymerization (Fig. 3H; Fig. S7A; Movie 5, 6). However, upon GTP addition, the disassembly of Shi rings resulted in the exposure of actin bundle sides, thus allowing robust branched actin polymerization from their sides (Fig. 3H; Fig. S7A; Movie 5, 6). Similarly, disassembly of human Dyn1 rings upon GTP addition also led to robust Arp2/3-mediated branched actin polymerization (Fig. S7B; Movie 7). Together, our data suggest that GTP hydrolysis driven disassembly of dynamin rings exposes the sides of the actin bundles to allow the generation of new short and branched actin filaments suitable for mechanical work. These branched filaments can be bundled by newly assembled dynamin rings to further increase the mechanical strength of the branched actin network to promote membrane protrusions.

To test whether GTP-induced disassembly of the Shi rings is required for cell-cell fusion, we performed rescue experiments using the G domain mutants in S2R+ cells co-expressing Sns, Eff-1 and dsRNA against the 5’ UTR of *shi*. Shi-Q35E, Shi-K137A and the ts2 mutant (at restrictive temperature) failed to rescue the cell fusion defect in Shi KD cells (Fig. 3I, J). In addition, overexpression of Shi-Q35E in wild-type S2R+ cells coexpressing Sns and Eff-1 caused strong dominant-negative effect on cell-cell fusion, whereas Shi-K137A had a weaker dominant-negative effect (Fig. 3I), consistent with the partial disassembly of Shi-K137A ring in the presence of GTP (Fig. 3B). Taken together, these results support a role for GTP hydrolysis-mediated dynamin ring disassembly in promoting cell-cell fusion.

### The stalk region and the PH domain are not specifically involved in dynamin-actin interactions

To further probe the mechanism of dynamin-actin bundling, we set out to map the specific actin-binding sites in dynamin. A previous study mapped the actin-binding domain of human Dyn1 to a region within the stalk, and showed that mutating several positively charged amino acids (Dyn1-K414/415/419/421/426E; known as Dyn1-AKE) significantly impaired Dyn1-actin interaction^29^. We made the corresponding Shi quadruple mutant, Shi-K410E, R411E, K417E, K422E (Shi-AKE), but found that the expression level of Shi-AKE was significantly lower (20.2% ± 15.7%) than wild-type Shi and most of the mutant protein was unstable during purification (Fig. S8A, B). Shi-AKE expression neither rescued the cell fusion nor endocytosis defect in Shi KD cells (Fig. S8C, D, E), and failed to cause a dominant-negative effect in cell fusion or endocytosis in wild-type cells (Fig. S8C, E). These results suggest that the Dyn1-AKE mutant likely affects Shi protein stability and assembly, rather than specifically blocking Shi-actin interactions.

The ability of dynamin to bind and assemble onto lipid nanotubes requires interactions with the PH domain^15^. Thus, we made a Shi mutant carrying a deletion of the PH domain (ShiΔPH) and tested its interaction with actin (Fig. 4A). High-speed and low-speed F-actin co-sedimentation assays and EM of negative stained samples revealed that ShiΔPH bound and bundled actin as effectively as wild-type Shi (Fig. 4B,C; Fig. S9A), but failed to form helices around lipid tubules as predicted (Fig. S9B). Similarly, a point mutation in the PH domain (Shi-I527A, corresponding to Dyn1-I533A), which is required to interact with PIP2, did not affect actin bundling (Fig. 3B). However, and unexpectedly, Shi-I527A failed to disassemble in the presence of GTP (Fig. 3B), suggesting again that the nature of dynamin-assembly with actin was structurally distinct from dynamin’s assembly around lipid templates. Consistent with a defect in disassembly, Shi-I527A failed to rescue the cell-cell fusion defect in Shi KD cells, and caused a dominant negative effect on cell-cell fusion in wild-type cells (Fig. 3I). These results suggest that actin filaments are not bundled in the center of the Shi rings, and that the PH domain is involved in conformational changes required to disassemble actin-associated Shi rings upon GTP hydrolysis.

**Fig. 4.**
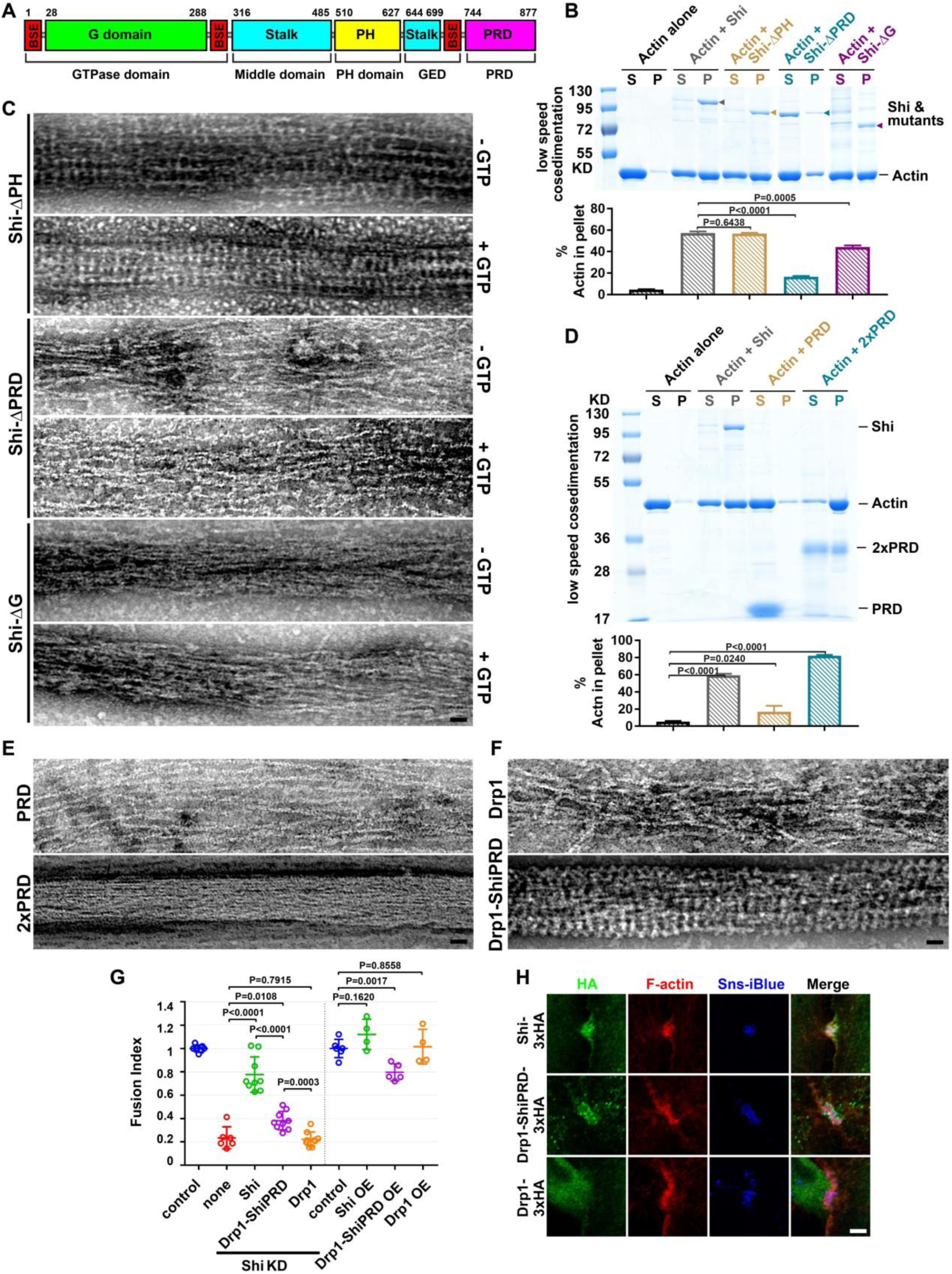
The PRD of Shibire mediates Shibire-actin binding. (A) Schematic diagram of the Shi domain structure. The amino acid numbers at the beginning and end of some domains are indicated. (B) The PRD and G domains, but not the PH domain, are involved in actin bundling. Actin-bundling activity of Shi and three deletion mutants (Shi-ΔPH, Shi-ΔPRD and Shi-ΔG) revealed by low speed co-sedimentation assay. Actin (3 μM) was incubated alone or with Shi (or Shi mutant) (0.5 μM). Assays were performed and quantified as in Fig. 2A. The Shi protein and mutants are indicated by arrowheads. Significance was determined by the two-tailed Student’s t-test. Values in the bar graph represent mean±SD (n = 3). Note that deleting PRD significantly compromised actin bundling. (C) Transmission electron micrographs of negatively stained actin filaments in the presence of Shi deletion mutants (Shi-ΔPH, Shi-ΔPRD and Shi-ΔG), in the absence or presence of GTP. Note that Shi-ΔPH bundled actin normally, but the Shi-ΔPH ring were not disassembled upon GTP addition; Shi-ΔPRD failed to bundle actin; and Shi-ΔG loosely bundled actin, but failed to form regularly spaced rings. Scale bar, 30 nm. (D) A tandem PRD has actin-bundling activity. Actin-bundling activity of single PRD or tandem PRD (2xPRD) revealed by low speed co-sedimentation assay. Actin (3 µM) was incubated alone or with 0.5 µM Shi, 1.5 µM PRD or 1 µM 2xPRD, respectively. Assays were performed and quantified as in Fig. 2A. Significance was determined by the two-tailed Student’s t-test. Values in the bar graph represent mean ± SD (n = 5). Note that 2xPRD exhibited a high actin-bundling activity. (E) Transmission electron micrographs of negatively stained actin filaments in the presence of PRD or 2xPRD. Note that PRD did not bundle actin, but 2xPRD induced tight actin bundles. Scale bar, 30 nm. (F) Transmission electron micrographs of negatively stained actin filaments in the presence of Drp1 or Drp1-ShiPRD. Note that Drp1 loosely bundled actin, but Drp1-ShiPRD induced tight actin bundles with regularly spaced Drp1-ShiPRD rings. Scale bar, 30 nm. (G) Quantification of the fusion index of S2R+ cells expressing Shi, Drp1-ShiPRD and Drp1. Note that in Sns-Eff-1-expressing Shi knockdown (KD) cells, Drp1-ShiPRD, but not Drp1, partially rescued the cell fusion defect. In Sns-Eff-1-expressing cell, overexpression (OE) of Drp1-ShiPRD, but not Drp1, caused a dominant negative effect in inhibiting cell fusion. All Shi and mutant constructs were tagged with 3xHA. The numbers of independent experiments performed: n = 7, 6, 9, 9, 9, 6, 4, 5 and 4 (left to right). The wide horizontal bars indicate the mean values. Significance was determined by the two-tailed Student’s t-test. (H) Drp1-ShiPRD, but not Drp1, is enriched at the fusogenic synapse of S2R+ cells. Sns/Eff-1-expressing S2R+ cells were labeled by anti-HA (green), phalloidin (F-actin; red) and Sns-iBlueberry (blue). The fusogenic synapses were marked by F-actin and Sns enrichment. Scale bar: 5 µm.

### The PRD of dynamin plays a primary role in mediating dynamin-actin interactions

Deleting the PRD domain (ShiΔPRD) significantly decreased Shi’s actin binding and bundling activities, shown by high-speed and low-speed co-sedimentation assays (Fig. 4B; Fig. S9A). Consistent with this, negative-stain EM revealed that ShiΔPRD failed to organize actin filaments in tight bundles and failed to form regularly spaced rings (Fig. 4C), despite forming normal helices around lipid tubules (Fig. S9B). Since PRD is best known for interacting with SH3 domains, we tested whether isolated SH3 domains would interfere with PRD-actin interaction. As shown in Fig. S9C, although SH3 domains compromised Shi-actin interaction, a significant portion of Shi remained in the pellet in the high-speed sedimentation assay, suggesting that the PRD can accommodate interactions with both SH3 domains and actin simultaneously (Fig. S9C). Indeed, most of the SH3-interacting PxxP motifs reside in the N-terminal half of the Shi PRD, whereas most of the conserved and positively charged amino acids reside in the C-terminal half of the PRD (Fig. S9D). The critical function of the PRD in actin binding is not specific for *Drosophila* Shi, since hDyn2ΔPRD also exhibited significantly decreased activities in actin binding and bundling (Fig. S4F, G). Interestingly, deleting the G domain (ShiΔG) also decreased Shi’s actin bundling activity, albeit to a lesser extent than deleting the PRD (Fig. 4B,C; Fig. S9A). Negative stain EM revealed that ShiΔG could bundle actin to some extent (likely due to the presence of the PRD) without forming regularly spaced rings (Fig. 4C). Taken together, these results suggest that the PRD is the primary actin-binding domain in Shi and that the G domain may also play a minor role in facilitating PRD in bundling actin and/or in enhancing ring assembly so as to position the PRDs for efficient actin binding.

To test whether the PRD itself has actin-binding activity, we purified the Shi PRD alone, as well as two PRDs linked in a tandem repeat (2xPRD). PRD alone exhibited a low actin-binding activity detected by high-speed F-actin co-sedimentation assay (Fig. S9E), and the PRD did not bundle actin as probed by low-speed F-actin co-sedimentation assay and negative-staining EM (Fig. 4D,E). However, 2xPRD showed a higher actin-binding activity (Fig. S9E) and organized tight actin bundles despite not forming regularly spaced rings like Shi (Fig. 4D, E). Therefore, the PRD binds actin and the tandem PRD serves as an artificial two-filament actin crosslinker. To further test the ability of PRD in actin bundling, we fused the Shi PRD domain to the C-terminus of *Drosophila* dynamin-related protein 1 (Drp1), which does not have a PRD and is normally required for mitochondria fission. Although the human Drp1 (hDrp1) has been shown to bundle actin by light microsocpy^39^, our negative stain EM revealed relatively loose actin bundles organized by either *Drosophila* Drp1 or hDrp1 without regularly spaced Drp1 bands (Fig. 4F; Fig. S10). Strikingly, however, the Drp1-ShiPRD fusion protein organized tight actin bundles with regularly spaced rings as Shi (Fig. 4F; Fig. S10), demonstrating a critical function of PRD in mediating the formation of tight actin bundles. Consistent with this, Drp1-ShiPRD, but not Drp1, partially rescued the fusion defect in Shi KD cells expressing Sns and Eff-1 (Fig. 4G). The partial rescue in vivo may be due to the requirement of a higher concentration of Drp1-ShiPRD than that of Shi to organize tight actin bundles (Fig. S10). Consistent with this, overexpressing (OE) Drp1-ShiPRD, which was enriched at the fusogenic synapse (Fig. 4H), exhibited a dominant negative effect in the fusion of wild-type cells, presumably by interfering endogenous Shi function as a less efficient actin-bundling protein (Fig. 4G). Taken together, these results demonstrate that the PRD is the primary actin-binding domain in Shi.

### Point mutations in the PRD differentially inhibit actin bundling and endocytosis

To gain further insight into the PRD-actin interaction, we made point mutations in the four highly conserved, positively charged residues in the Shi PRD, R804D, R829D, R846D, and R853D (Shi-4RD) (Fig. S9D), and tested the interactions between the dynamin mutants with negatively charged actin. While R804D resides in a PxxP motif and R829 is two amino acids away from a PxxP motif, the two other mutations, R846D and R853D, did not overlap with any PxxP motifs (Fig. S9D). The single mutants (Shi-R804D and Shi-R829D) and the double mutant (Shi-R846/853D) did not affect Shi-actin interaction shown by low/high-speed F-actin co-sedimentation assays and negative-staining EM (Fig. 5A, B, C). However, the quadruple mutant (Shi-R804/829/846/853D or Shi-4RD) exhibited a significant decrease in actin binding and bundling, demonstrating the specificity of the PRD-actin interaction (Fig. 5A, B, C). Furthermore, Shi-4RD, but not the single or double mutants, failed to rescue the fusion defect in Sns/Eff-1-expressing Shi KD cells (Fig. 5D, E), despite their normal colocalization with the F-actin-enriched foci at the fusogenic synapse, likely mediated by PxxP-SH3 domain interactions (Fig. 5F; Fig. S11A). Although Shi-4RD was unable to rescue fusion, this mutant could significantly, albeit partially, rescue the endocytosis defect in Shi KD cells (Fig. 5G, H; Fig. S11B). These data, together with our inability to detect endocytic structures at the fusion synapse even when fission is blocked (Fig. 1I), suggest that dynamin’s function in actin bundling is uncoupled from its role in endocytosis. The incomplete rescue of endocytosis by Shi-4RD supports a potential involvement of dynamin-mediated actin bundling in endocytosis.

**Fig. 5.**
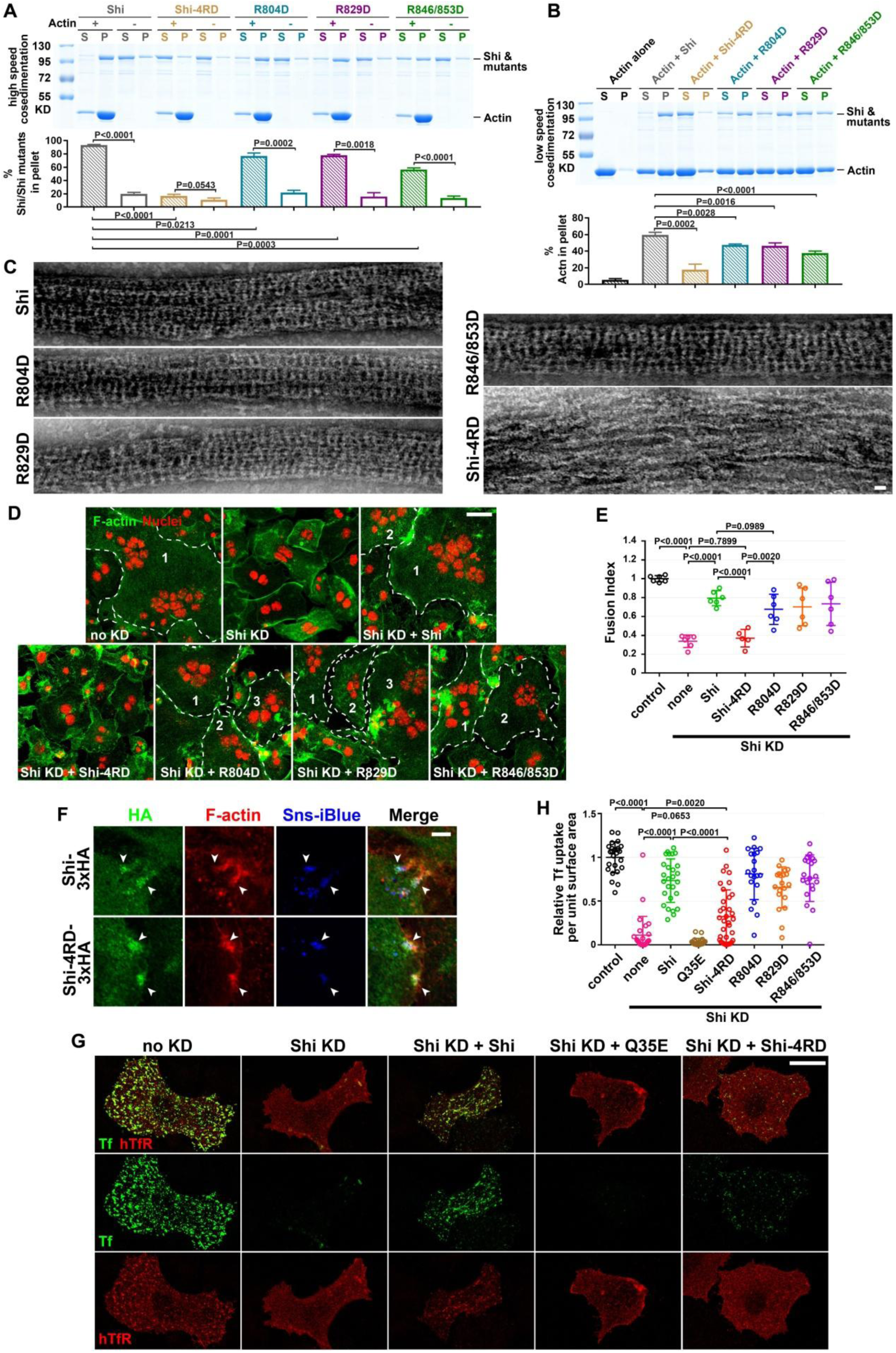
Four highly conserved arginine residues are essential for Shibire-actin interaction and cell-cell fusion. (A) High speed co-sedimentation assays reveal the actin-binding activity of Shi and its PRD domain mutants (Shi-4RD, R804D, R829D and R846/853D). Shi or PRD-mutants (0.5 µM) were incubated with or without actin (3 µM), and supernatants (S) and pellets (P) were monitored by SDS-PAGE after centrifugation at 50,000*g*. The percentage of Shi or the PRD-mutants in the pellet vs. total was quantified. Significance was determined by the two-tailed Student’s t-test. Values in the bar graph represent mean±SD (n = 3). (B) Low speed co-sedimentation assays reveal the actin-bundling activity of Shi and its PRD domain mutants. Actin (3 µM) was incubated alone or with Shi or PRD-mutants (0.5 µM). Assays were performed and quantified as in Fig. 2A. Significance was determined by the two-tailed Student’s t-test. Values in the bar graph represent mean±SD (n = 4). (C) Transmission electron micrographs of negatively stained actin filaments in the presence of Shi or PRD-mutants. Note that Shi-4RD failed to bundle actin. Scale bar, 30 nm. (D) Shi-4RD failed to rescue the fusion defect caused by Shi knockdown in S2R+ cells. Sns/Eff-1-expresssing S2R+ cells fused to form multinucleated (≥ 3 nuclei) syncytia (outline and numbered). Cells were labeled with phalloidin (F-actin; green) and Hoechst 33342 (nuclei; red), and fused cells were identified as in Fig. 1D. Shi was knocked down (KD) with dsRNA targeting Shi 5’UTR and inhibited cell fusion. Note that expressing the PRD single and double mutants, but not Shi-4RD, rescued the fusion defect. Scale bar: 20 µm. (E) Quantification of the fusion index of S2R+ cells for each condition in (D). The numbers of independent experiments performed: n = 6, 6, 6, 5, 6, 6 and 6 (left to right). The wide horizontal bars indicate the mean values. Significance was determined by the two-tailed Student’s t-test. (F) Shi-4RD is enriched at the fusogenic synapse of S2R+ cells. Sns/Eff-1-expressing S2R+ cells were labeled by anti-HA (green), phalloidin (F-actin; red) and Sns-iBlueberry (blue). The fusogenic synapses were marked by F-actin and Sns enrichment. Scale bar: 5 µm. (G) Shi-4RD partially rescued endocytosis in Shi knockdown (KD) S2R+ cells. Pulse-chase transferrin (Tf) uptake assays were performed with Shi or Shi mutants in Shi KD S2R+ cells expressing human transferrin receptor (hTfR). Cells were labeled with Tf-Alexa488 (green) and anti-hTfR (red). Note that Shi KD led to a failure in Tf uptake, which was partially rescued by the expression of Shi-4RD but not Q35E (indicated by internal Tf vesicles). Scale bar: 20 µm. (H) Quantification of Tf uptake assays shown in (G) and Fig. S11B, measured by the number of Tf vesicles per unit surface area of the cell. The numbers of cells examined: n = 25, 28, 27, 22, 35, 20, 20 and 20 (left to right). The wide horizontal bars indicate the mean values. Significance was determined by the two-tailed Student’s t-test.

### Multiple actin filaments are locked to the outer rim of the dynamin helix in each actin bundle

That the externally oriented PRD is the primary site for actin binding suggests that actin filaments are likely to be attached to the outer rim of the dynamin ring. We next applied negative stain and cryo-ET to further explore the nature of dynamin-actin interactions. In the negative stain EM experiments with a low concentration of Shi (0.125 μM), actin filaments were occasionally not bundled together by dynamin at the ends. Such “split ends” allowed us to discern ∼8 actin filaments within an actin bundle (Fig. 6A, top panel). In addition, in some middle segments of an actin bundle where Shi rings were absent, we detected actin filaments running along the outer rim of the Shi rings (Fig. 6A, bottom panel). These data strongly support our model that the actin filaments bind to the outer edges of the Shi rings.

**Fig. 6.**
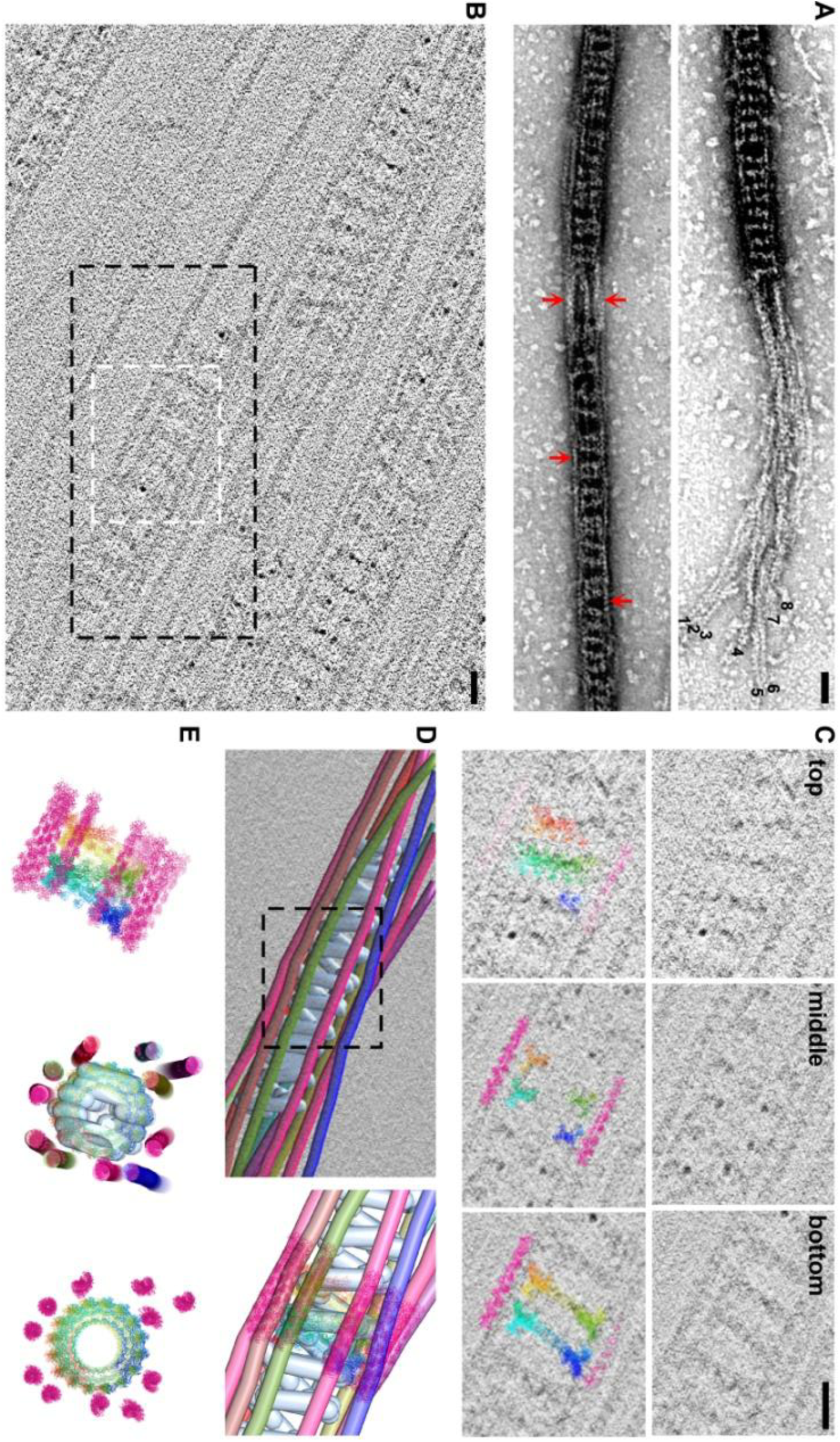
Dynamin forms a helix that engages multiple actin filaments at the outer rim. (A) Electron micrographs of negatively stained Shi-mediated actin bundles. Eight actin filaments can be observed at the “split” end of the actin bundle (top panel). Actin filaments (red arrows) are located at the outer rim of the Shi rings. Scale bar, 30 nm. (B) Cryo-ET section illustrating several hDyn1-mediated actin bundles. Sections through the white-boxed area are shown in (C). Black box outlines region selected for model building in imod shown in (D). Scale bar, 20 nm. (C) Sections through the cryo-tomogram with the molecular model manually docked into the tomogram. The top section reveals a right-handed tilt of hDyn1, the middle shows the T-shaped structures, and the bottom section shows a left-handed tilt of hDyn1, consistent with a helical polymer. Scale bar, 20 nm. (D) Model of a hDyn1-mediated actin bundle manually generated in imod with the hDyn1 helix colored light blue and actin filaments on the outside rainbow colored. A molecular model of hDyn1 and actin docked into the imod model is shown on the right (boxed area on the left). (E) Side view (left panel) and top views (middle and right panels) of a molecular model of the hDyn1-mediated actin bundle. The middle panel shows the top view of the molecular model docked in the imod model. Note that not all of actin filaments are associated with the hDyn1 helix on this particular cross section.

To achieve a higher resolution view of dynamin-mediated actin bundles, we performed cryo-electron tomography (cryo-ET) using hDyn1 (or Shi) with actin. Cryo-ET revealed that hDyn1 assembled as a helix surrounded by actin filaments (Fig. 6B-E; Fig. S12A, B; Movie 8-11). The hDyn1 helices were evident by sectioning through the tomogram and observing that the hDyn1 bands on the top of the bundle had a right-handed tilt, whereas the corresponding hDyn1 bands on the bottom of the bundle exhibited a left-handed tilt (Fig. 6C; Fig. S12A; Movie 9). The middle section of the bundle contained T-shaped structures, similar to images of dynamin decorated lipid tubes^13^, with the top of the T near the actin filaments that were positioned at the outer rim of the dynamin helix (Fig. 6C; Fig. S12A; Movie 9). Based on the outer diameter (42 nm) and the pitch (16.8 nm) of the hDyn1 helix observed by cryo-ET, we made a model of the hDyn1 helix and positioned actin filaments based on the tomogram (Fig. 6D, E; Fig. S12B). Since the length per turn (L) of the hDyn1 helix was 133 nm, [L = (C^2 + H^2) ^(0.5); C = circumference, H = height or pitch], and the distance between hDyn1 dimers is 8.3 nm based on the high-resolution cryo-EM map of hDyn1^40^, we determined that there were ∼16 hDyn1 dimers per turn (133 nm divided by 8.3 nm) in the actin bundle, which correlated with the helical parameters of 22.5-degree twist and 10.5 Å rise between dimers. Segmentation of a hDyn1 helix and its associated actin filaments revealed 11 actin filaments associated with the outer rim of this hDyn1 helix (Fig. 6D, E; Fig. S12B; Movie 9-11), corresponding to a partially occupied hDyn1 helix in which 11 of the 16 PRDs per turn engaged with actin filaments. Cryo-ET also revealed that *Drosophila* Shi assembled as a helix surrounded by actin filaments (Fig. S12C, D; Movie 12). Based on the outer diameter (32 nm) and the pitch (15.1 nm) of Shi, we estimated that there are ∼12 Shi dimers per turn in the actin bundle. The 8-filament bundle shown in Fig. 6A therefore is a partially occupied Shi helix. We conclude from these results that each dynamin-mediated actin bundle consists of a centrally localized dynamin helix and multiple actin filaments attached to the outer rim of a dynamin helix.

## Discussion

Our data demonstrate that dynamin oligomerizes to regulate actin bundling as it does in regulating endocytosis. However, the mechanisms by which dynamin helices and rings interact with different substrates, such as lipid tubules vs. actin filaments, are distinct. While the lipid tubule is wrapped in the center of a dynamin helix via its interaction with the PH domain, the actin filaments are locked to the outer rim of dynamin rings via their specific interactions primarily with the PRD (Fig. S13A). Our data demonstrate that the PRD itself can directly bind and bundle actin without other linker proteins, in contrast to previous studies suggesting that the PRD interacts with actin filaments through a linker protein, cortactin^22,31,32^. In vivo studies have shown that the homozygous *Drosophila cortactin* mutant is viable and phenotypically normal^41^, and that double knockout of cortactin and its related protein HS1 in mouse platelets does not affect the formation of podosomes or platelet function^42^. In addition, pharmacological inhibition or genetic depletion of cortactin did not affect actin comet tail formation or *Listeria* motility in *Listeria*-infected mammalian cells^43,44^, whereas dynamin depletion significantly reduces both^24,25,44^. Thus, cortactin does not appear to have an essential role in dynamin-mediated actin bundling. Although the PRD is best known for its role in dynamin localization via its interaction with the SH3 domain-containing proteins, the PRD can accommodate SH3 and actin interactions simultaneously, likely due to the largely separated SH3-interacting PxxP motifs and the positively charged amino acids involved in actin binding. The identification of the positively charged amino acids in the PRD allows us to uncouple dynamin’s function in actin bundling vs. fission of lipid tubules. In this regard, Shi-4RD abolishes dynamin’s activity to bundle actin and promote cell fusion, but only partially affected its activity in supporting endocytosis. Thus, the dynamin-actin interactions are essential for actin bundling, and only facilitates, but is not required for, endocytosis.

Our study establishes dynamin as a unique F-actin-bundling protein. Most actin bundling factors, such as *α*-actinin, fascin, fimbrin, filamin, myosin II, and spectrin are two-filament crosslinkers^45-47^. They form linear or V-shaped structures (can be monomer, dimer or heterotetramer), each of which has two actin-binding sites located at the two opposite ends of the molecule and thus crosslinking two actin filaments like a bridge. Dynamin, on the other hand, appears to be a multi-filament bundler. Dynamin does this by forming a helical structure composed of either 16 dimers (hDyn1) or 12 (Shi) dimers per helical turn, allowing the binding of a maximal number of 16 (hDyn1) or 12 (Shi) actin filaments to a single helix (Fig. S13A). Due to the tilt of the dynamin dimers along the long axis of the actin bundle, each actin filament is bound by two PRDs from two different dimers in each turn of a dynamin helix. By interacting with additional PRDs along the dynamin helix, the multi-filament actin bundle is further stabilized. Moreover, the multi-filament single bundles can align in parallel to form super bundles, likely mediated by trans-bundle PRD-actin interactions. The ability to mediate the formation of multi-filament single bundles, as well as super bundles, makes dynamin a unique and efficient actin-bundling protein.

Our data strongly support a critical function of dynamin in mechanical force generation. By forming single bundles and super bundles, dynamin increases the mechanical strength of the actin filaments. Moreover, the actin-stimulated GTP hydrolysis allows dynamic disassembly of dynamin helices, freeing the sides of the actin bundles and promoting Arp2/3-meidated branched actin polymerization. The short, branched actin filaments may join the pre-existing, partially occupied actin bundles or form new bundles de novo via actin-PRD interactions, thus further increasing the mechanical strength of the actin filaments (Fig. 13B). Thus, by equilibrating between helix assembly and disassembly as a function of GTP hydrolysis, dynamin gradually increases the mechanical strength of the actin network. As a consequence, the podosome-like structure at the fusogenic synapse is able to propel membrane protrusions (Fig. 13C) and the comet tails are able to propel bacteria movement. We speculate that the mechanisms underlying dynamin-actin interaction uncovered in this study will be generally applicable to other cellular processes involving dynamin-mediated actin cytoskeletal rearrangements.

---

**Fig. S1.**
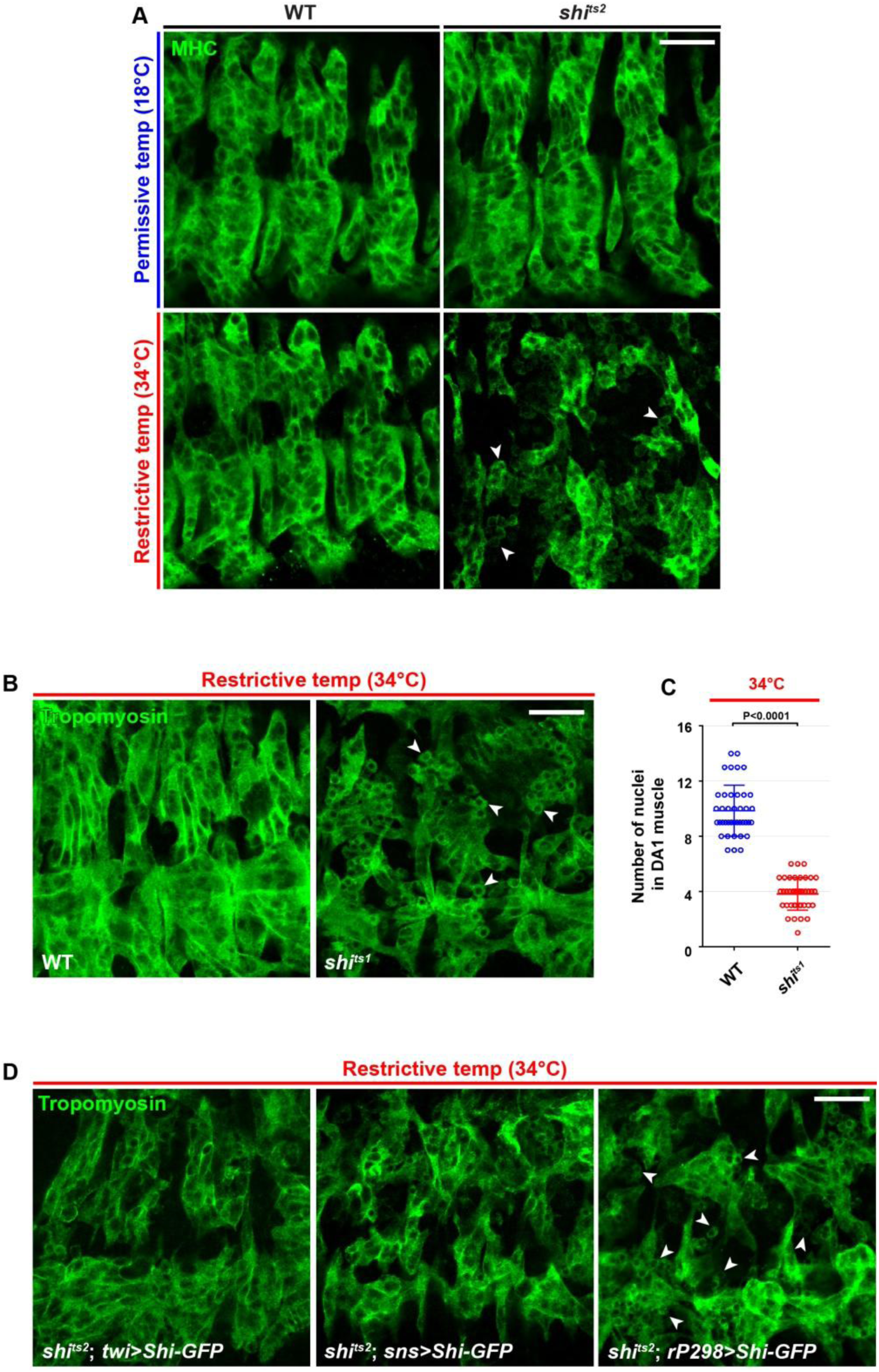
Shi is required for myoblast fusion. (A) Myoblast fusion defect in *shi^ts2^*mutant embryo. The somatic muscles in stage 15 wild-type (WT) and *shi^ts2^* mutant embryos were labeled by anti-muscle myosin heavy chain (MHC) antibody. Note the presence of unfused mononucleated myoblasts in *shi^ts2^* mutant embryo at the restrictive temperature (arrowheads). Scale bar: 20 µm. (B) Myoblast fusion defects in *shi^ts1^*mutant embryo. The somatic muscles in stage 15 wild-type (WT) and *shi^ts1^* mutant embryos were labeled by anti-Tropomyoson antibody. Note the presence of unfused mononucleated myoblasts in *shi^ts1^*mutant embryo at the restrictive temperature (arrowheads). Scale bar: 20 µm. (C) Quantification of the fusion index of genotypes shown in (B). The numbers of DA1 muscles analyzed were 40 for WT and 40 for *shi^ts1^*. The wide horizontal bars indicate the mean values. Significance was determined by the two-tailed Student’s t-test. (D) Phenotypic rescue of myoblast fusion in *shi^ts2^* mutant embryos at restrictive temperature. Note that expression of Shi-GFP in all muscles (*twi-GAL4*) or FCMs (*sns-GAL4*), but not in founder cells (*rP298-GAL4*), restore myoblast fusion in *shi^ts2^* embryos at the restrictive temperature. Unfused mononucleated myoblasts are indicated by arrowheads (see quantification in Fig. 1B). Scale bar: 20 µm.

**Fig. S2.**
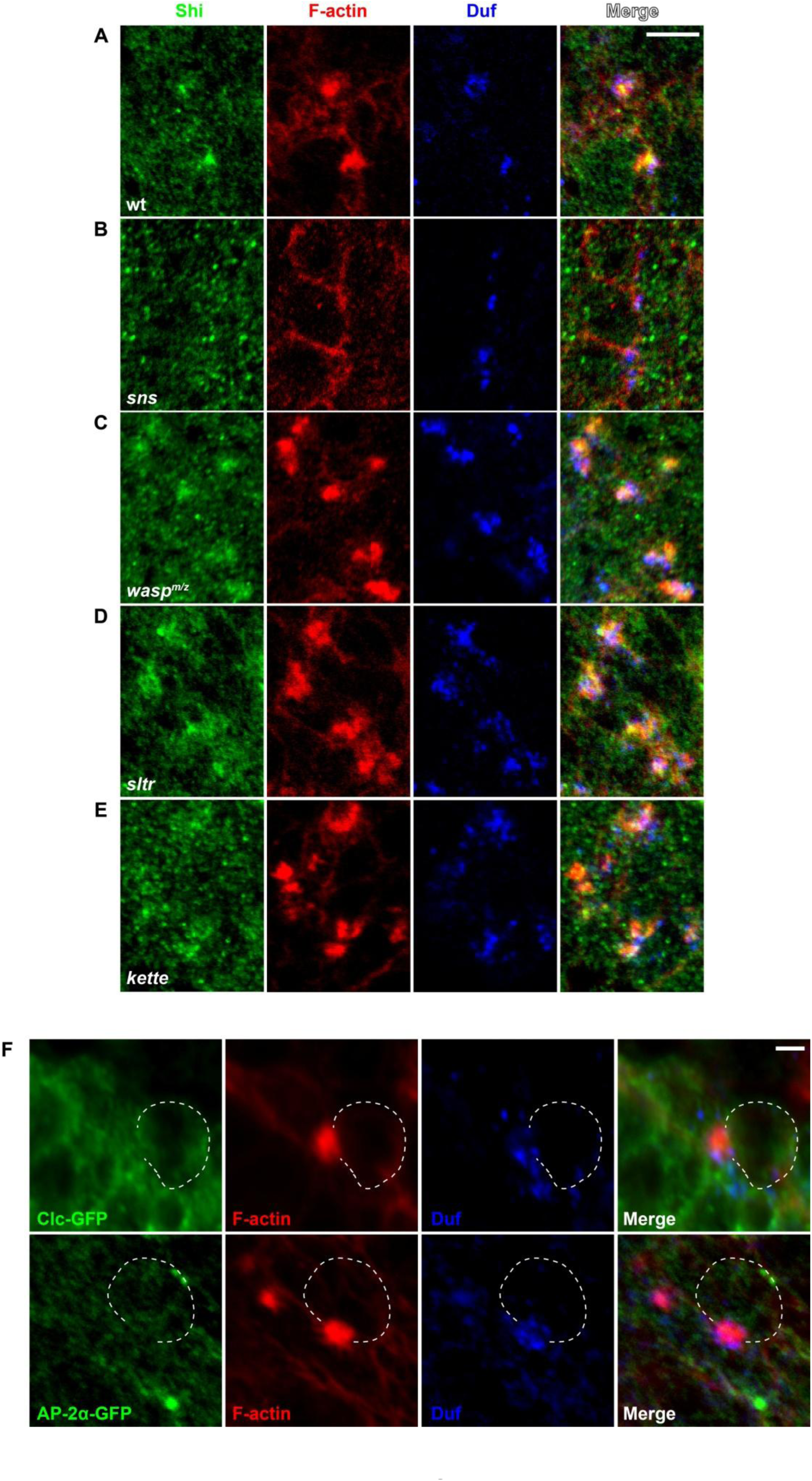
Localization of endogenous Shi and the endocytic proteins at the fusogenic synapse. (A-E) The recruitment of Shi to the fusogenic synapse is dependent on the FCM-specific cell adhesion molecule, but not on actin polymerization regulators. Stage 14 embryos of various genotypes were labeled with anti-Shi (endogenous Shi; green), phalloidin (F-actin; red), and anti-dumbfounded (Duf; founder cell-specific adhesion molecule enriched at the fusogenic synapse; blue). Shi and F-actin were enriched at the fusogenic synapse in WT (A), *wasp*^m/z^ (C), *sltr* (D), and *kette* (E) mutant embryos, but not in *sns* mutant embryo (B). Scale bar: 5 µm. (F) Clathrin light chain (Clc) (top panels) and the α subunit of the AP-2 clathrin adaptor complex (AP-2α) (bottom panels) are not enriched at the fusogenic synapse. Stage 14 embryos expressing Clc-GFP or AP-2α-GFP were labeled with phalloidin (F-actin; red) and anti-Duf (blue). The FCM is outlined. Note that F-actin, but not Clc-GFP (or AP-2α), was enriched at the fusogenic synapse. Scale bar: 2 µm.

**Fig. S3.**
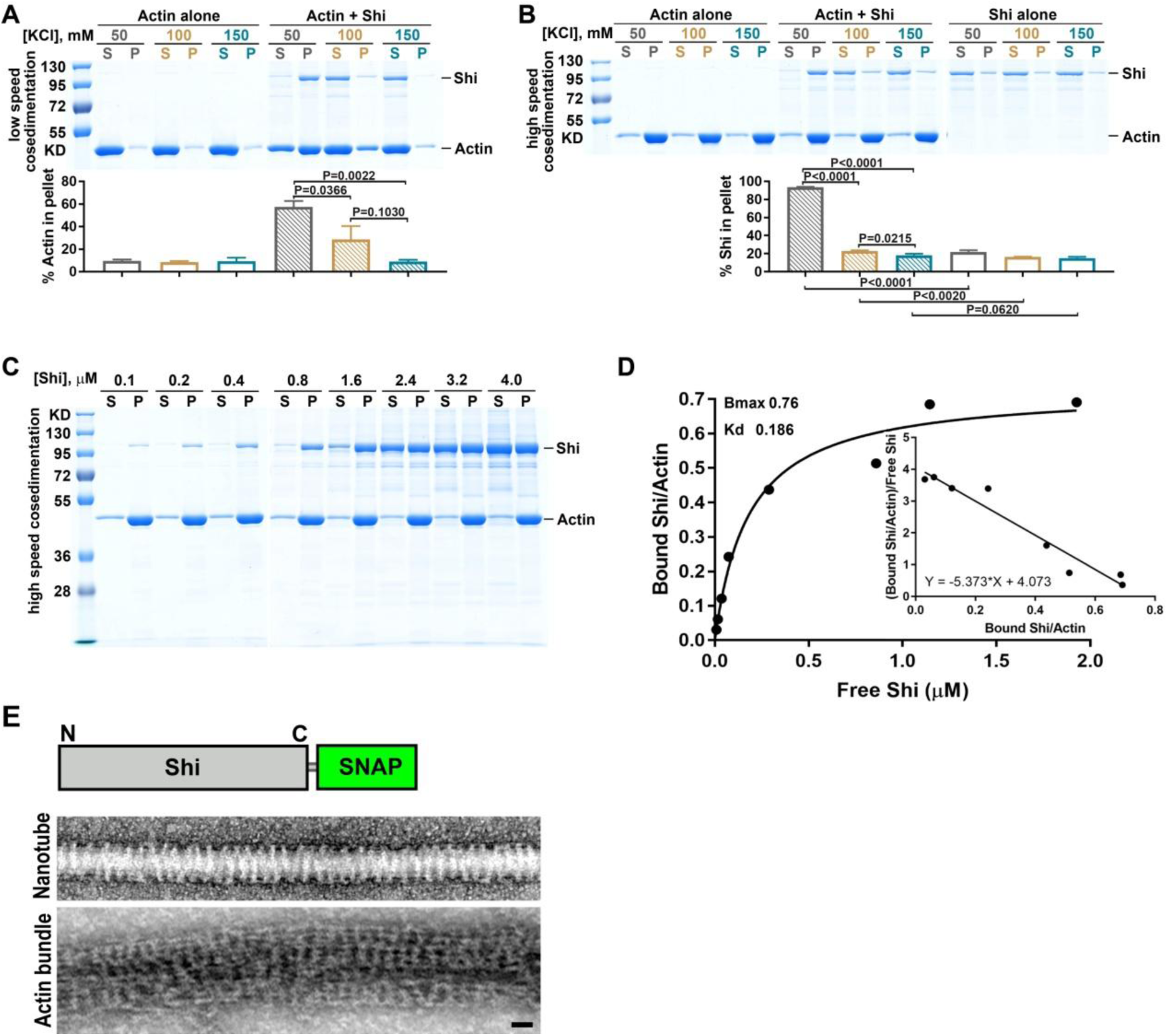
Shi bundles actin filaments and forms helices around lipid nanotubes. (A) Shi bundles actin filaments at a low salt condition. Low speed co-sedimentation assays were performed to assess the actin-bundling activity of Shi at different salt (KCl) concentrations as indicated. 3 μM actin was incubated alone or with 0.5 μM Shi. Assays were performed and quantified as in Fig. 2A. Values in the bar graph represent mean±SD (n=3). Significance was determined by two-tailed Student’s *t*-test. (B) Shi binds actin filaments at a low salt condition. High speed co-sedimentation assays were performed to assess the actin-binding activity of Shi at different salt (KCl) concentrations as indicated. 0.5 μM Shi was incubated alone or with 3 μM actin. Assays were performed and quantified as in Fig. 5A. Values in the bar graph represent mean±SD (n=3). Significance was determined by two-tailed Student’s *t*-test. (C) High speed co-sedimentation assays were performed with 3 μM actin and increasing concentrations of Shi to calculate the stoichiometry of binding and dissociation equilibrium constant. Supernatants (S) and pellets (P) were monitored by SDS-PAGE after centrifugation at 50,000*g*. (D) Analysis of Shi-actin binding shown in (C). The concentration of bound Shi was plotted against the concentration of free Shi and fitted to a hyperbolic function (one site binding). Scatchard analysis of Shi-actin binding was performed to calculate the stoichiometry of binding and dissociation equilibrium constant. (E) Shi-SNAP is functional. A schematic diagram of Shi with a C-terminal SNAP-tag is shown at the top. Bottom panels show transmission electron micrographs of negatively stained lipid nanotube wrapped with Shi-SNAP helices, and an actin super bundle organized by Shi-SNAP. Scale bar, 30 nm.

**Fig. S4.**
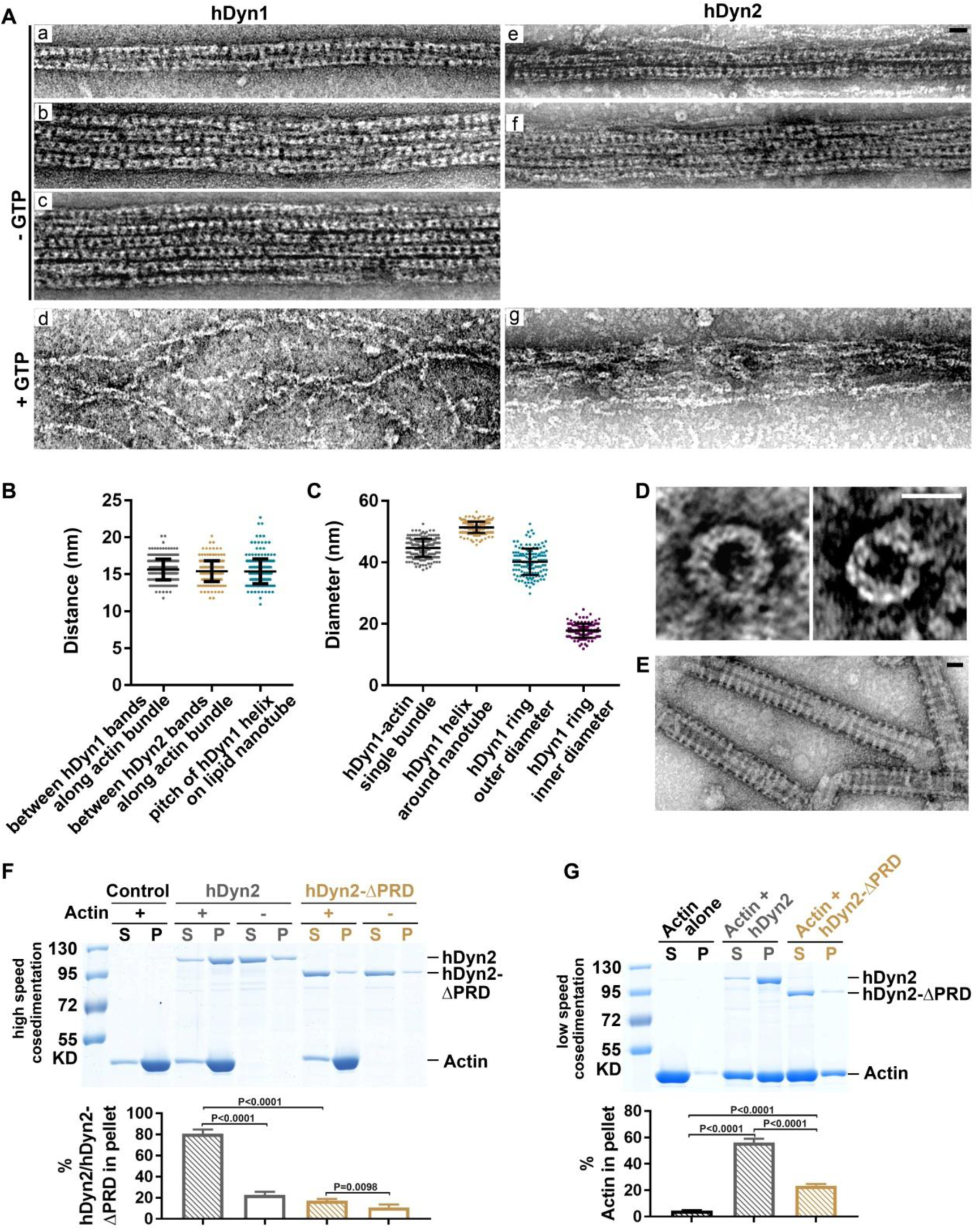
Human dynamin 1 and 2 bundle actin filaments. (A) Transmission electron micrographs of negatively stained actin filaments organized by hDyn1 (a-d) or hDyn2 (e-g) in the absence or presence of GTP. Note that both hDyn1 and hDyn2 formed rings to bundle actin to form single (a, e), double (b, f), and triple (c) bundles. GTP addition disassembled the dynamin rings and the actin bundles (d, g). Scale bar, 30 nm. (B) Dot plot of the distance between hDyn1 or hDyn2 rings alone the actin bundle, and the pitch of hDyn1 helices on lipid nanotube shown in (E). Values in the bar graph represent mean±SD, *n* = 307, 245 and 252, respectively. (C) Dot plot of the diameters of hDyn1-actin single bundle and hDyn1 helix around nanotube shown in (E), as well as the outer and inner diameters of hDyn1 ring shown in (D). Values in the bar graph represent mean±SD, n = 220, 226, 132 and 131, respectively. (D) Transmission electron micrographs of negatively stained hDyn1 rings. Scale bar, 30 nm. (E) Transmission electron micrographs of negatively stained lipid nanotubes wrapped in hDyn1 helices. Scale bar, 30 nm. (F) High speed co-sedimentation assays reveal the actin-binding activity of hDyn2 and hDyn2-ΔPRD. 0.5 μM hDyn2 (or hDyn2-ΔPRD) was incubated alone or with 3 μM actin. Assays were performed and quantified as in Fig. 5A. Values in the bar graph represent mean±SD (n=4). Significance was determined by two-tailed Student’s *t*-test. Note the significant decrease of actin-binding activity of hDyn2-ΔPRD compared to that of hDyn2. (G) Low speed co-sedimentation assays reveal the actin-bundling activities of hDyn2 and hDyn2-ΔPRD. 3 μM actin was incubated alone or with 0.5 μM hDyn2 (or hDyn2-ΔPRD). Assays were performed and quantified as in Fig. 2A. Values in the bar graph represent mean±SD (n=3). Significance was determined by two-tailed Student’s *t*-test. Note the significant decrease of actin-bundling activity of hDyn2-ΔPRD compared to that of hDyn2.

**Fig. S5.**
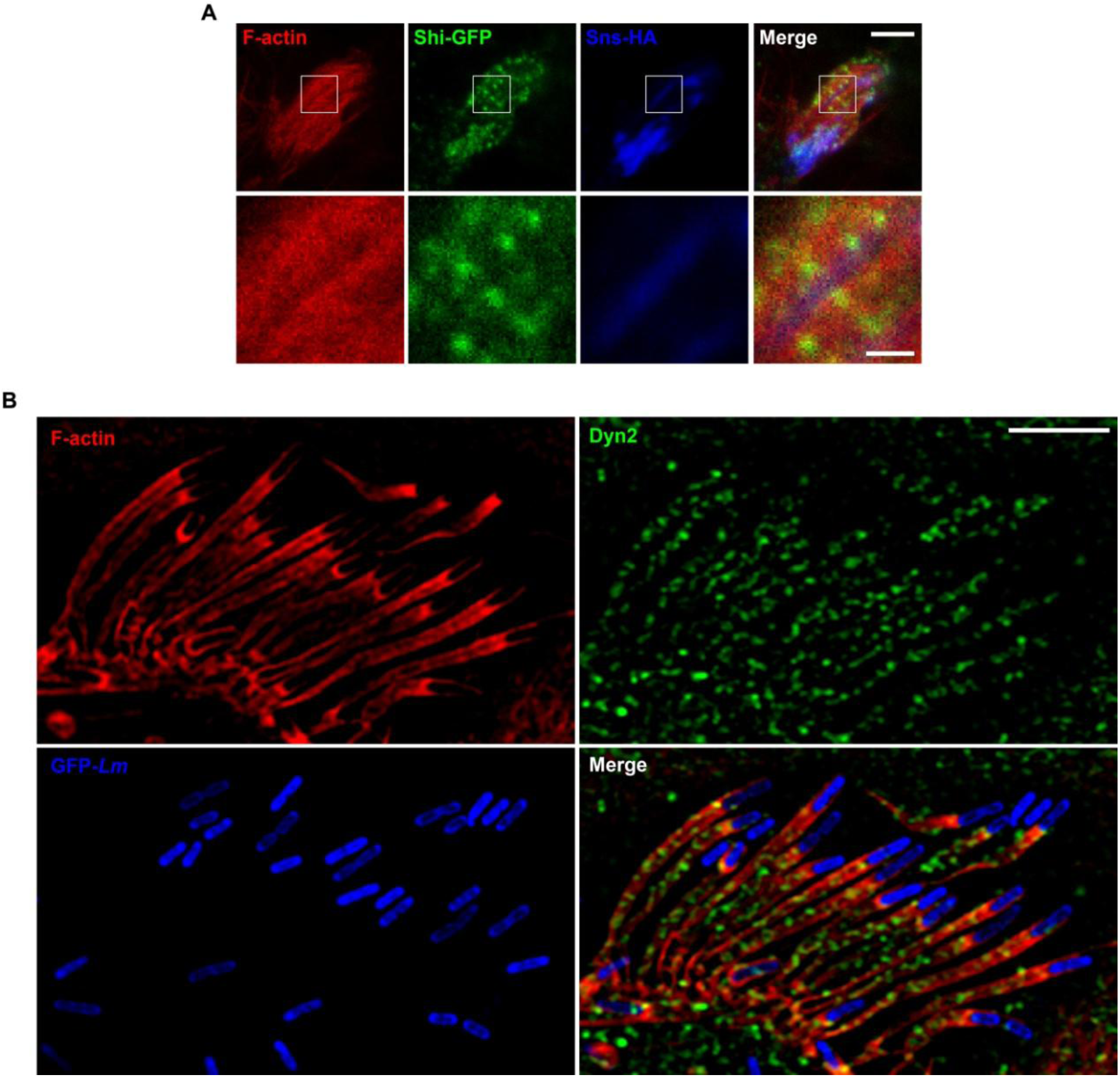
Dynamin colocalization with actin bundles at fusogenic synapses and actin comet tails. (A) Shi localization on actin bundles at the fusogenic synapse revealed by stimulated emission depletion microscopy (STED). S2R+ cells co-expressing Sns-HA (blue), Eff-1-FLAG and Shi-GFP (green) were labeled with phalloidin (F-actin; red). Boxed areas are enlarged in the insets. Note the Shi punctae and bands along the actin bundles. Scale bars, 2 μm (main panels) or 500 nm (insets). (B) hDyn2 localization on actin comet tails in *Listeria-*infected HeLa cell revealed by structured illumination microscopy (SIM). Over two dozen *Listeria* (*Listeria*-GFP; blue) were captured protruding out of a host cell, each propelled by an actin comet tail. Note the hDyn2 punctae and bands (green) along each comet tail labeled with phalloidin (red). Scale bars: 4 µm.

**Fig. S6.**
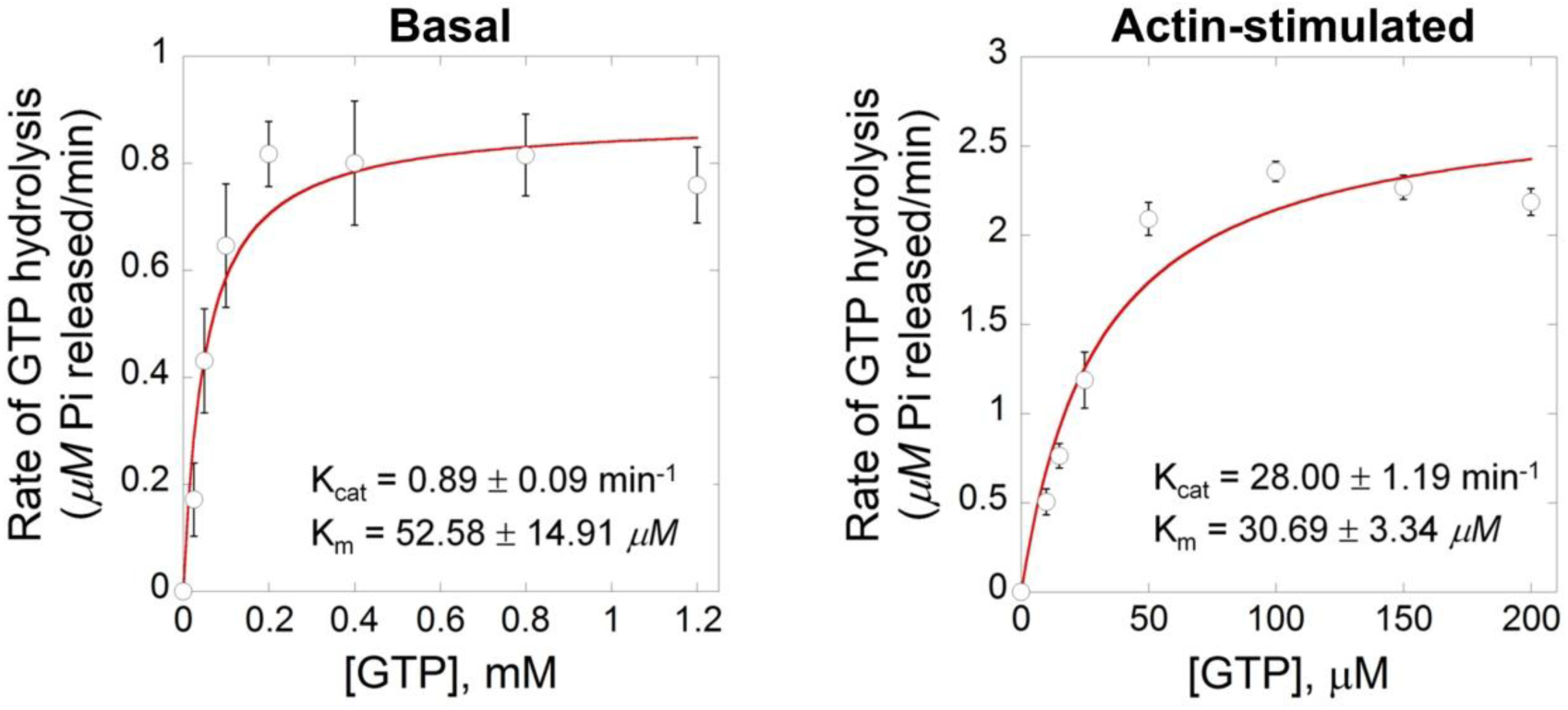
Actin bundling stimulated the GTP hydrolysis rate of Shi. GTP hydrolysis rates of Shi at basal state (left) and actin-stimulated state (right) were plotted against the initial concentrations of GTP to calculate the Michaelis-Menten constants K_cat_ and K_m_. Each data point represents mean±SD, n = 3 and 4, respectively. Note the significantly enhanced K_cat_ in the presence of actin filaments.

**Fig. S7.**
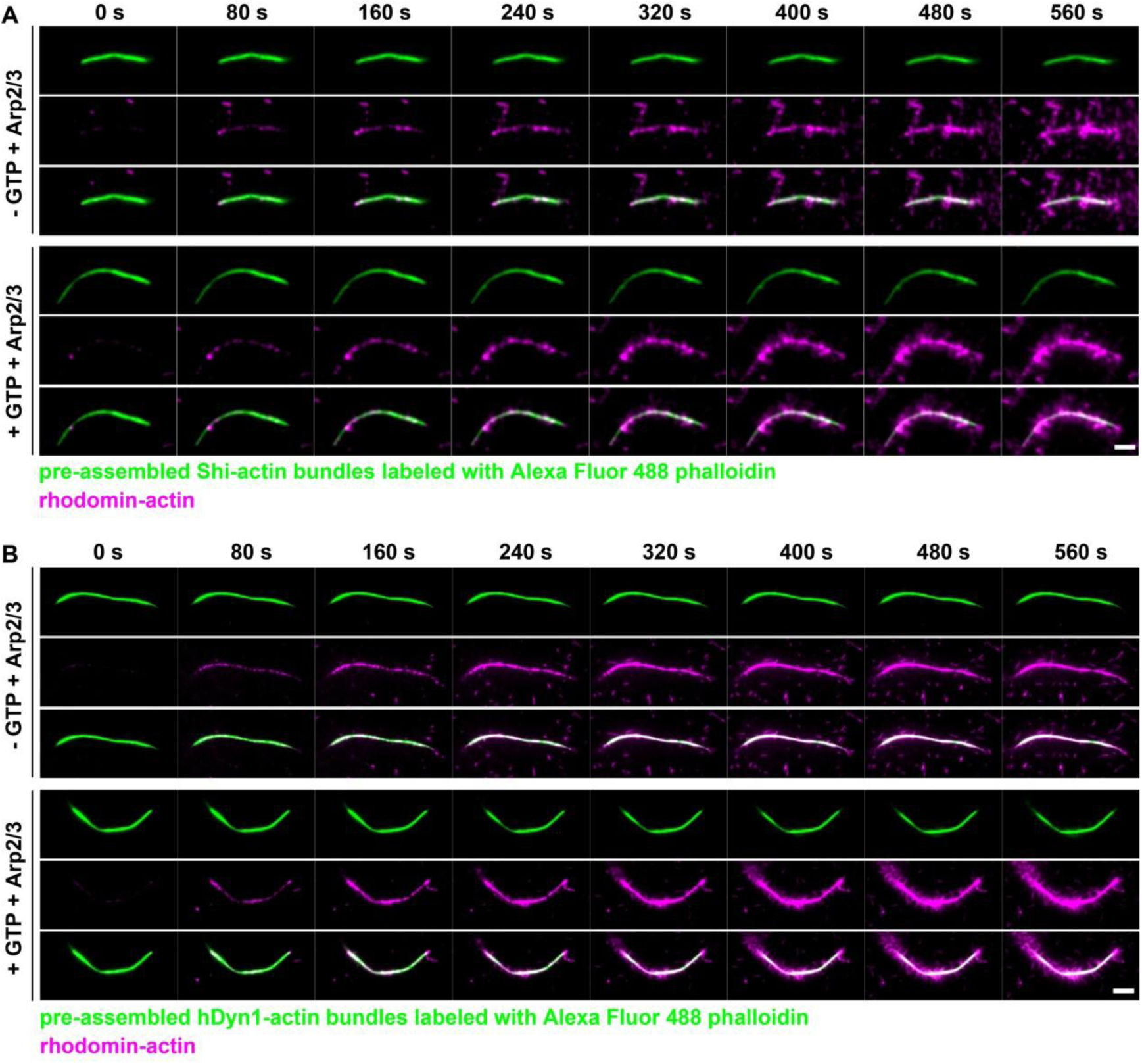
GTP addition leads to increased Arp2/3-mediated branched actin polymerization on dynamin-actin bundles. (A) Time-lapse stills of TIRF images showing Arp2/3-mediated branched actin polymerization on Shi-actin bundles in the absence or presence of GTP (see Movie 6). 2 μM Shi was first incubated with 1 μM Alexa Fluor 488 phalloidin-labeled actin (green) to generate actin bundles. Subsequently, Arp2/3, VCA and rhodamine-actin (magenta) were added to the Shi-actin bundles to start branched actin polymerization without (top panels) or with (bottom panels) 1 mM GTP. Note the enhanced amount of branched actin filaments emanating from the actin bundles with GTP compared to without GTP. Scale bar, 5 μm. (B) Time-lapse stills of TIRF images showing Arp2/3-mediated branched actin polymerization on hDyn1-actin bundles in the absence or presence of GTP (see Movie 7). 2 μM hDyn1 was first incubated with 1 μM Alexa Fluor 488 phalloidin-labeled actin (green) to generate actin bundles, followed by a wash with TIRF buffer without (top panels) or with (bottom panels) 0.01 mM GTP. Subsequently, Arp2/3, VCA and rhodamine-actin (magenta) were added to start branched actin polymerization. Scale bar, 5 μm.

**Fig. S8.**
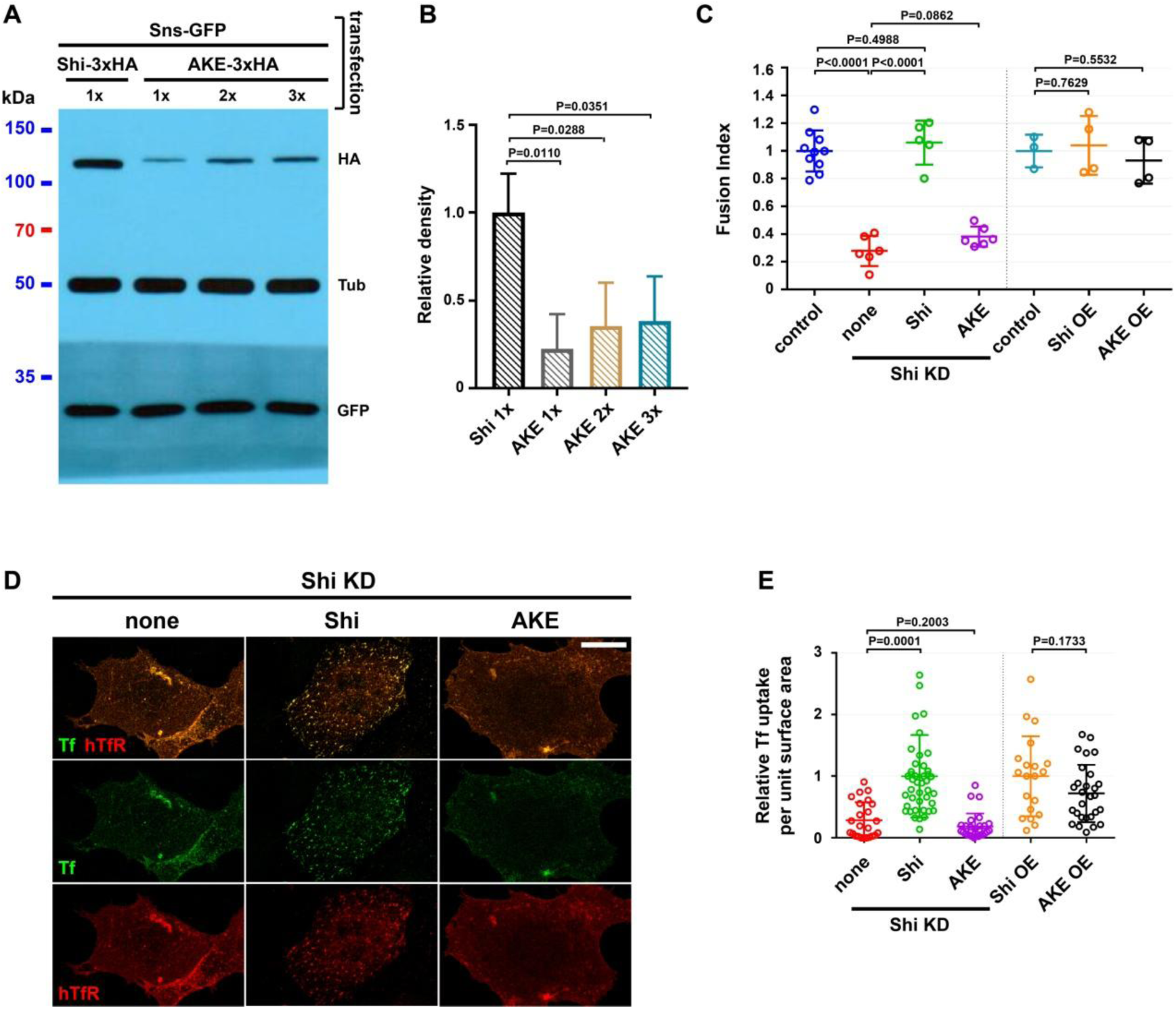
The stalk domain mutant Shi-AKE fails to rescue cell fusion or endocytosis defects in Shi knockdown cells. (A) Shi-AKE is less stable than Shi shown by western blot analysis. Note the reduced AKE-3xHA protein level compared to that of Shi-3xHA, even when 3x plasmid was transfected. Sns-GFP and α-Tubulin were used as transfection and loading controls, respectively. (B) Quantification of the western blot shown in (A). Values in the bar graph represent mean±SD (n=3). Significance was determined by two-tailed Student’s *t*-test. Note the significantly lower expression of the AKE mutant than wild-type Shi. (C) Quantification of the fusion index of Sns/Eff-1-expressing S2R+ cells. Note that unlike Shi-3xHA, AKE-3xHA was unable to rescue the fusion defect in Shi knockdown (KD) S2R+ cells. Overexpressing (OE) AKE-3xHA in S2R+ cells did not cause a dominant negative effect in cell fusion. The numbers of independent experiments performed: n = 10, 6, 5, 6, 3, 4 and 4 (left to right). The wide horizontal bars indicate the mean values. Significance was determined by the two-tailed Student’s t-test. (D) Pulse-chase transferrin (Tf) uptake assays were performed with wild-type Shi or Shi-AKE mutant in Shi KD S2R+ cells. Cells were labeled with Tf-Alexa488 (green) and anti-human Tf receptor (hTfR) (red). Note that Shi KD led to a failure in Tf uptake, which was rescued by Shi-3xHA, but not Shi-AKE-3xHA, expression (indicated by internal Tf vesicles). Scale bar: 20 µm. (E) Quantification of Tf uptake assays shown in (D), measured by the number of Tf vesicles per unit surface area of the cell. The numbers of cells examined: n = 24, 43, 28, 20 and 28 (left to right). The wide horizontal bars indicate the mean values. Significance was determined by the two-tailed Student’s t-test.

**Fig. S9.**
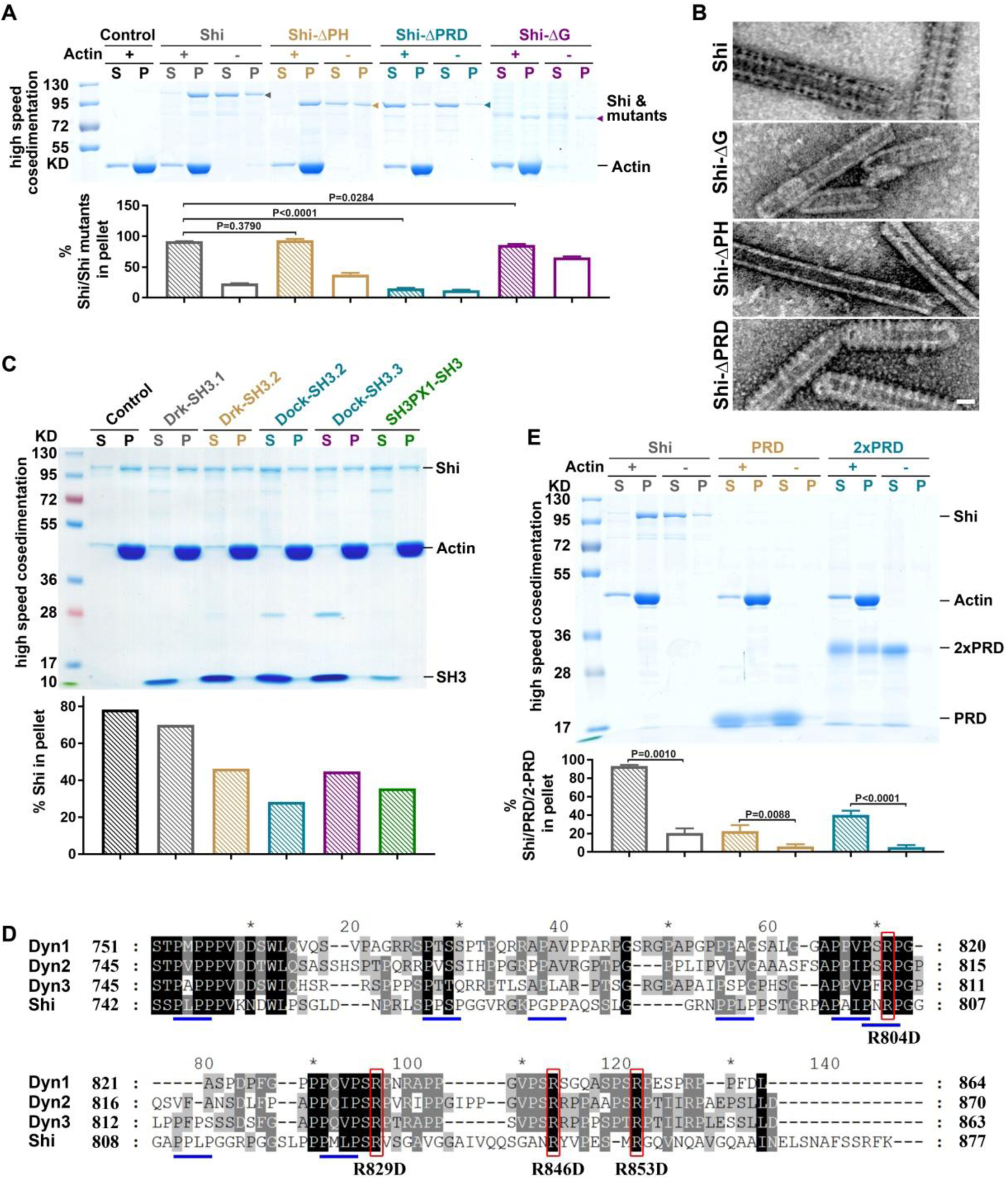
The PRD domain of Shi mediates Shi-actin binding. (A) High speed co-sedimentation assays reveal the actin-binding activity of Shi and three deletion mutants (Shi-ΔPH, Shi-ΔPRD and Shi-ΔG). 0.5 μM Shi (or Shi mutant) was incubated alone or with 3 μM actin. Assays were performed and quantified as in Fig. 5A. The Shi protein and mutants are indicated by arrowheads. Values in the bar graph represent mean±SD (n=3). Significance was determined by two-tailed Student’s *t*-test. Note the significant decrease of actin-binding activity of Shi-ΔPRD compared to that of Shi. (B) Transmission electron micrographs of negatively stained lipid nanotubes that were incubated with Shi or Shi deletion mutants, Shi-ΔPH, Shi-ΔG and Shi-ΔPRD. Scale bar, 30 nm. (C) SH3 domains interfere with, but do not block, Shi-actin interaction. High speed co-sedimentation assays reveal the effects of SH3 domains, Drk-SH3.1 (the first SH3 domain of Drk), Drk-SH3.2 (the second SH3 domain of Drk), Dock-SH3.2 (the second SH3 domain of Dock), Dock-SH3.3 (the third SH3 domain of Dock), and SH3PX1-SH3 (the SH3 domain of SH3PX1), to Shi-actin interaction. 6 μM actin was incubated with 0.5 μM Shi in the absence or presence of a SH3 domain. Assays were performed and quantified as in Fig. 5A. (D) Amino acid sequence alignment of the PRDs from human Dyn1 (NP_004399.2), human Dyn2 (NP_001005360.1), human Dyn3 (NP_056384.2), and *Drosophila* Shi (NP_727910.1). Red boxes indicate highly conserved arginine residues in the PRD. Blue underlines indicate PxxP motifs in Shi. (E) High speed co-sedimentation assays reveal the actin-binding activity of single PRD or two tandem PRDs (2xPRD). 0.5 μM Shi, 1.5 μM PRD or 1 μM 2xPRD was incubated alone or with 3 μM actin. Assays were performed and quantified as in Fig. 5A. Values in the bar graph represent mean±SD (n=4). Significance was determined by two-tailed Student’s *t*-test. Note that both PRD and 2xPRD exhibited actin-binding activity.

**Fig. S10.**
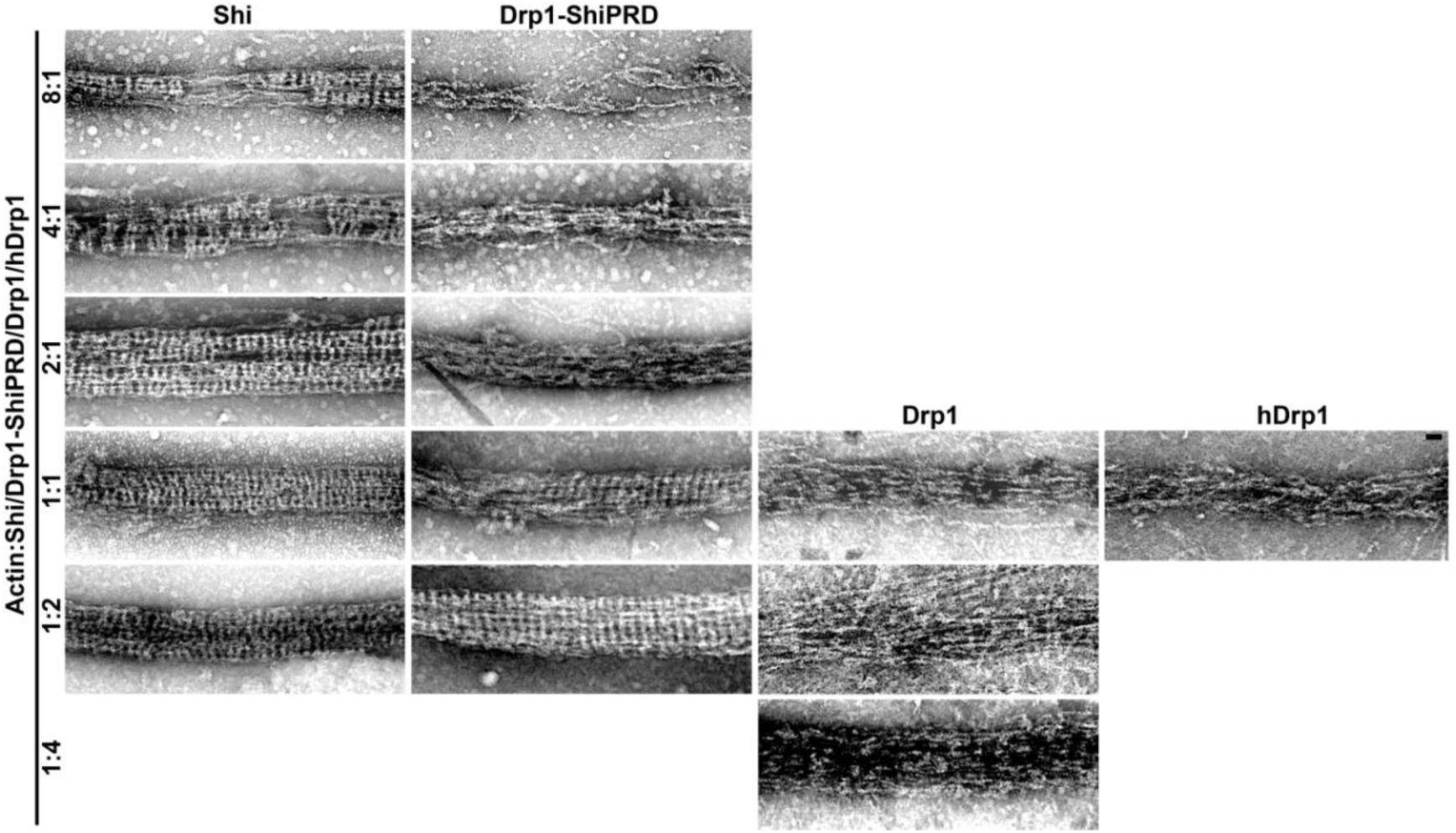
Drp1-ShiPRD bundles actin at a higher concentration than Shi. Transmission electron micrographs of negatively stained actin filaments in the presence of *Drosophila* Shi, Drp1-ShiPRD, Drp1, and human Drp1 (hDrp1) at different ratio of actin to Shi/Drp1-ShiPRD/Drp1/hDrp1 indicated on the left. Note that Shi formed rings along actin bundles when actin:Shi = 8:1, whereas Drp1-ShiPRD started to form rings along actin bundles when actin:Shi = 1:1. Note that neither *Drosophila* nor hDrp1 formed rings along actin bundles. Scale bar, 30 nm.

**Fig. S11.**
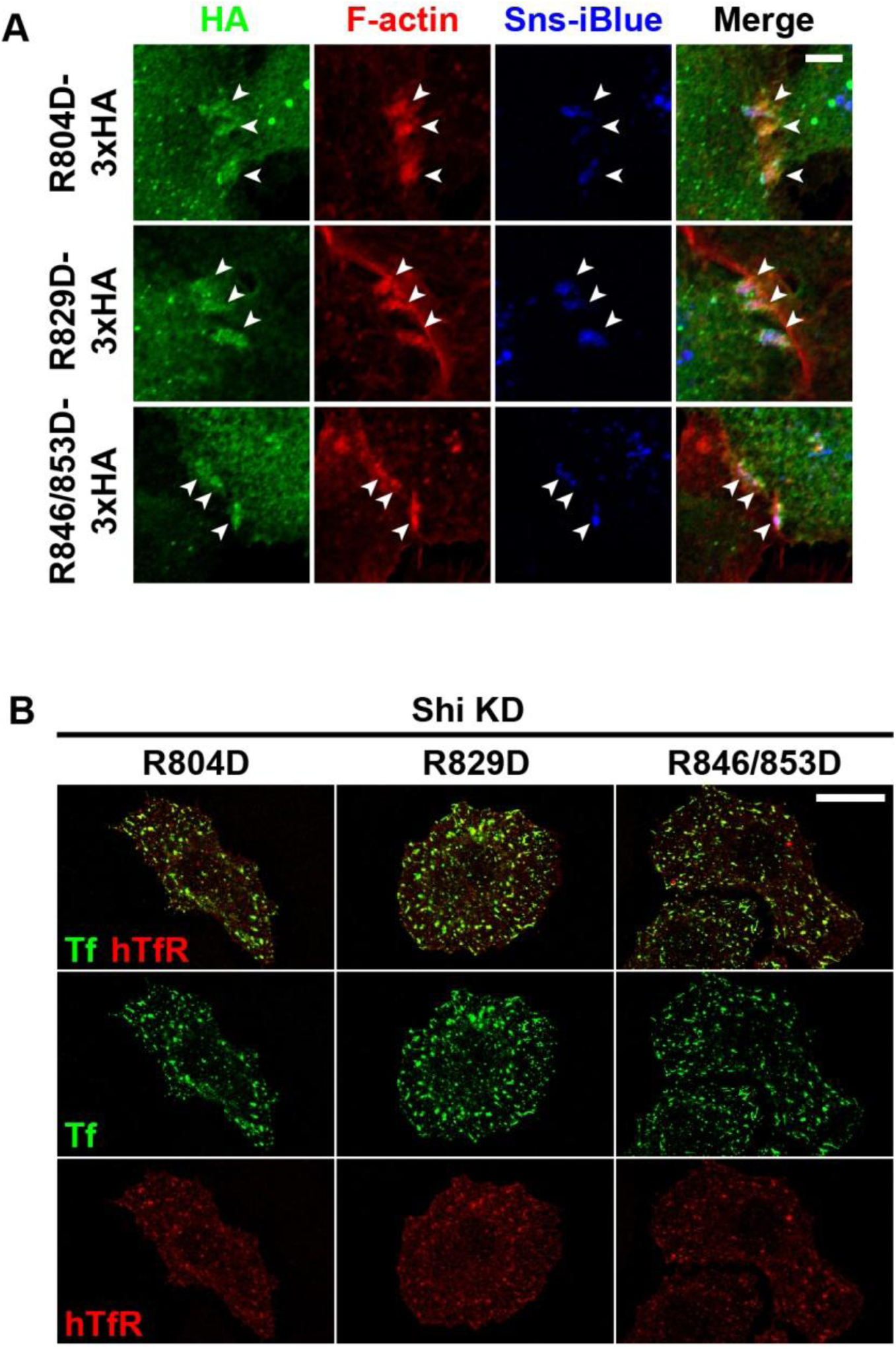
Single and double Shi PRD domain mutants are enriched at the fusogenic synapse and rescue the endocytosis defect in Shi knockdown cells. (A) Shi PRD domain mutants (R804D, R829D, and R846/853D) are enriched at the fusogenic synapse. Sns/Eff-1-expressing S2R+ cells were labeled by anti-HA (green), phalloidin (F-actin; red) and Sns-iBlueberry (blue). The fusogenic synapses were marked by F-actin and Sns enrichment. Scale bar: 5 µm. (B) Shi PRD domain mutants (R804D, R829D, and R846/853D) rescued endocytosis in Shi knockdown (KD) S2R+ cells. Pulse-chase transferrin (Tf) uptake assays were performed with Shi mutants in Shi KD S2R+ cells. Cells were labeled with Tf-Alexa488 (green) and anti-human Tf receptor (hTfR) (red). Note that the Shi mutants rescued the Tf uptake defect caused by Shi KD. Scale bar: 20 µm.

**Fig. S12.**
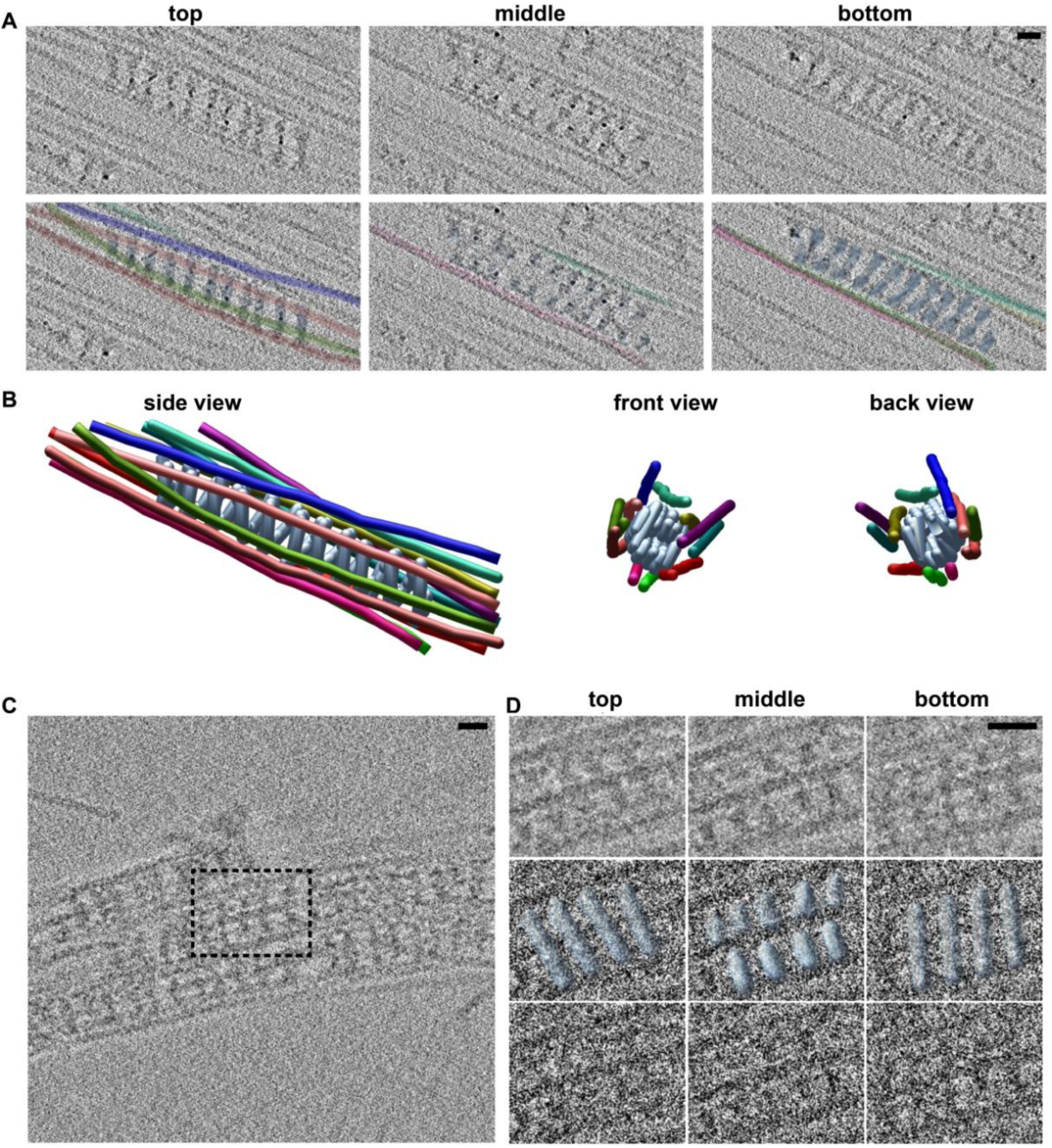
Cryo-ET of hDyn1- and Shi-mediated actin bundles. (A) Top, middle and bottom sections of cryo-ET showing actin filaments bundled by a hDyn1 helix. The top panels are sections of the tomogram and the bottom panels are the same sections with modelled hDyn1 (sky blue) and actin (other colors). Scale bar, 20 nm. (B) Tomogram-derived model of actin filaments bundled by a hDyn1 helix showing the side, front, and back views. Colors are the same as in (A). (C) Cryo-ET section illustrating several Shi-mediated actin bundles. Box outlines the region selected for modeling of Shi helix in (D). Scale bar, 20 nm. (D) Cryo-ET of Shi-mediated actin bundles reveals that Shi assembles as helices surrounded by actin filaments. Top, middle, and bottom views of the boxed region in (C) are consistent with a Shi helical polymer (colored blue). The top panel is the boxed area binned for ease of visualization. The middle panel has the modelled Shi helix docked onto the unbinned tomogram. The bottom panel is a view of the unbinned tomogram. Scale bar, 20 nm.

**Fig. S13.**
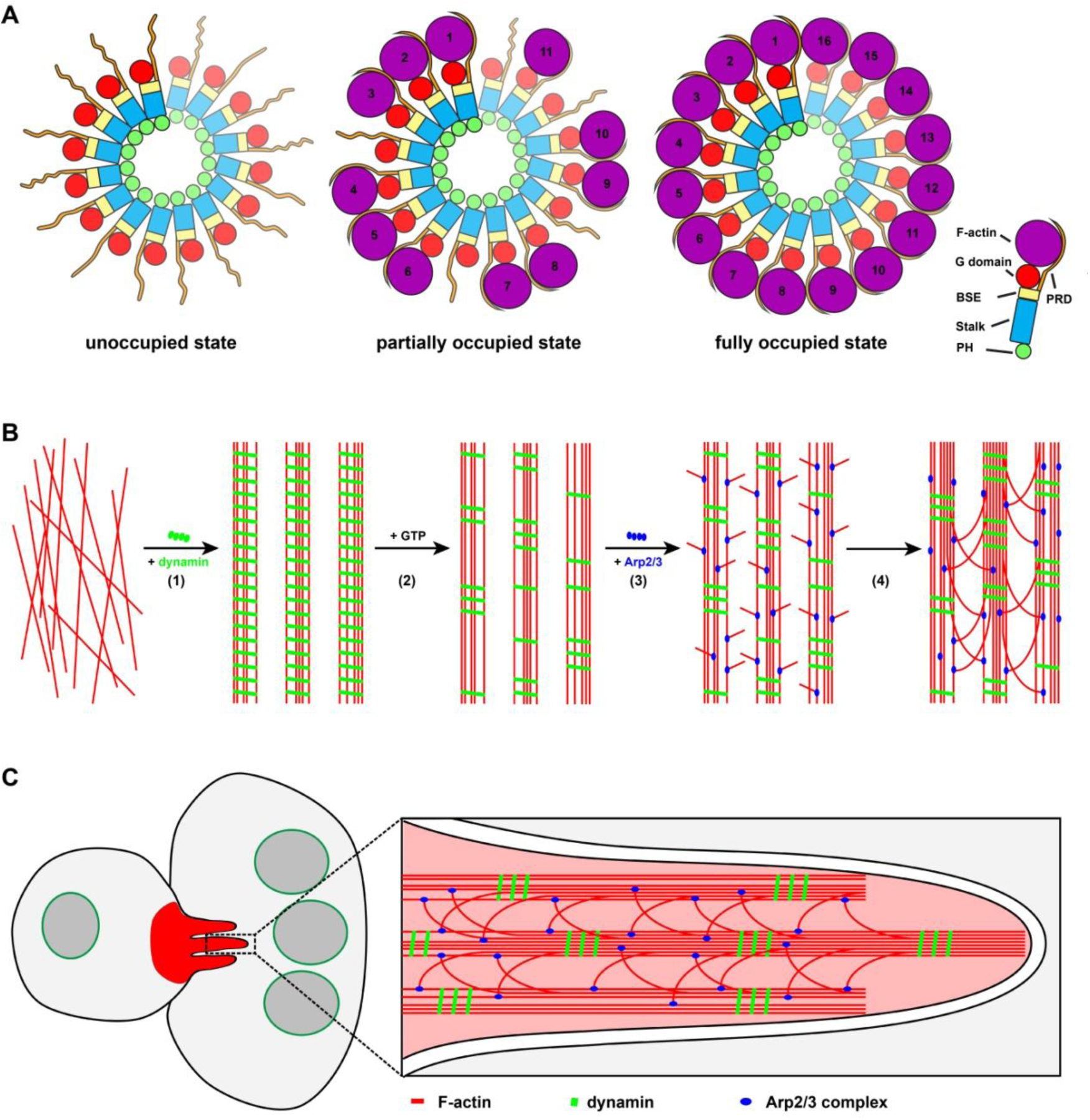
Models describing the mechanisms of dynamin-actin interaction. (A) Dynamin bundles multiple actin filaments. Schematic diagrams of single rungs of a dynamin (hDyn1 shown here) helix in three actin filament-(un)occupied states. Each hDyn1 rung contains 16 dimers, and only one monomer of each dimer is shown. The monomers more distant from the plane are depicted smaller and dimmer. In the unoccupied state (actin filament are absent), the PRDs extend outward from the dynamin helix and are unstructured. In the partially occupied state, some, but not all, PRDs lock actin filaments to the outer rim of dynamin. In the fully occupied state, all 16 PRDs have engaged actin filaments. (B) GTP hydrolysis of dynamin enhances Arp2/3-mediated branched actin polymerization. (1) In the absence of GTP, dynamin assembles into long helices that bundle actin filaments in vitro (red lines). (2) In the presence of GTP, GTP hydrolysis drives dynamin disassembly creating gaps in the dynamin helices and exposing regions of the bundled actin filaments. (3) Arp2/3 binds to the exposed sides of the actin bundles and initiates branched actin polymerization. (4) Branched actin filaments are further bundled by newly formed dynamin helices. (C) Dynamin promotes membrane protrusions at the fusogenic synapse. Dynamin bundles actin filaments to make them mechanically suitable for generating membrane protrusions. *In vivo* due to the presence of cytosolic GTP, dynamin undergoes dynamic cycles of assembly and disassembly. Assembled dynamin helices bundle actin filaments, while dynamin disassembly directs Arp2/3 complex-mediated actin polymerization to generate shorter branched actin filaments. The newly generated branched actin filaments may join partially occupied bundles and connect multiple bundles together. These dynamic events increase the mechanical stiffness of actin network and promote the formation of membrane protrusions.

**Supplementary Movie 1. Shi is dynamically associated with the F-actin focus at the fusogenic synapse.**

Time-lapse imaging of muscle cells expressing Shi-GFP (green) and Actin-mRFP (red) in a wild-type stage 14 embryo. The accumulation and dissolution of Shi coincided with that of the F-actin foci. 36 fusogenic synapses were imaged with similar results.

**Supplementary Movie 2. Shi bundles preassembled actin filaments.**

Time-lapse TIRF imaging of Shi-mediated bundling of preassembled actin filaments. Left, actin alone; right, actin and Shi. The TIRF assay was performed on supported lipid bilayers. Shi was added into the reaction mix of preassembled actin filaments to final concentrations of 10 nM (Shi) and 100 nM (actin). Three independent experiments were performed with similar results.

**Supplementary Movie 3. Shibire bundles growing actin filaments.**

Time-lapse TIRF imaging of Shi-mediated actin bundling during polymerization. The TIRF assay was performed in mPeg-Silane-coated flow chambers. To initiate the experiment, 1.5 μM G-actin and 1 μM Shi were mixed for imaging. Three independent experiments were performed with similar results.

**Supplementary Movie 4. GTP hydrolysis induces dynamic disassembly of Shi rings.**

FRAP of Shi-SNAP-Surface 488 along an actin bundle in the absence (left panel) or presence (right panel) of GTP. A segment of Shi-actin bundle was photobleached and the fluorescence recovery was monitored live. Note that the fluorescence was recovered in the presence, but not absence, of GTP.

**Supplementary Movie 5. GTP hydrolysis promotes dynamic Arp2/3-mediated branched actin polymerization on Shi-mediated actin bundles.**

Time-lapse TIRF imaging of Arp2/3-mediated branched actin polymerization on Shi-actin bundles in the absence (top panel) or presence (bottom panel) of GTP. Shi-actin bundles were generated by incubating unlabeled actin with Shi-SNAP-Surface 488 (green). Arp2/3, VCA and rhodamine-actin

(magenta) were added to the Shi-actin bundles to start branched actin polymerization without or with 1 mM GTP. Note the disassembly of Shi from the actin bundle upon GTP addition, and the numerous branched actin filaments emanating from the same actin bundle. 24 (or 14) Shi-actin bundles with (or without) GTP were imaged with similar results.

**Supplementary Movie 6. GTP addition leads to increased Arp2/3-mediated branched actin polymerization on Shi-actin bundles.**

Time-lapse TIRF imaging of Arp2/3-mediated branched actin polymerization on Shi-actin bundles in the absence (top panel) or presence (bottom panel) of GTP. Shi was first incubated with Alexa Fluor 488 phalloidin-labeled actin (green) to generate actin bundles. Subsequently, Arp2/3, VCA and rhodamine-actin (magenta) were added to the Shi-actin bundles to start branched actin polymerization without or with 1 mM GTP. Note the enhanced amount of branched actin filaments emanating from the actin bundles with GTP compared to without GTP. 20 (or 8) Shi-actin bundles with (or without) GTP were imaged with similar results.

**Supplementary Movie 7. GTP addition leads to increased Arp2/3-mediated branched actin polymerization on hDyn1-actin bundles.**

Time-lapse TIRF imaging of Arp2/3-mediated branched actin polymerization on hDyn1-actin bundles in the absence (top panel) or presence (bottom panel) of GTP. hDyn1 was first incubated with Alexa Fluor 488 phalloidin-labeled actin (green) to generate actin bundles, followed by a wash with TIRF buffer without or with 0.01 mM GTP. Subsequently, Arp2/3, VCA and rhodamine-actin (magenta) were added to start branched actin polymerization. Note the enhanced amount of branched actin filaments emanating from the actin bundles with GTP compared to without GTP. 22 (or 18) hDyn1-mediated actin bundles with (or without) GTP were imaged with similar results.

**Supplementary Movie 8. Cryo-electron tomogram of actin filaments incubated with hDyn1.**

The video shows z-sections of cryo-ET generated from a tilt series collected on cryo grids containing actin filaments and hDyn1. Scale bar, 10 nm.

**Supplementary Movie 9. Modelling hDyn1-actin interaction based on cryo-electron tomography.**

This movie includes: (1) z-sections of a cryo-electron tomogram; and (2) segmentation of actin filaments and hDyn1 by points in order to develop a model of actin-hDyn1 interaction. The hDyn1 helix is sky blue and actin filaments are in other colors. Scale bar, 10 nm.

**Supplementary Movie 10. Modelling actin filament interaction with hDyn1 based on cryo-electron tomography.**

This movie includes: (1) z-sections of a cryo-electron tomogram; (2) modelling in of the hDyn1 helix; (3) modelling of the actin filaments; (4) modelling of the hDyn1 helix and actin filaments; (5) model turning full 360° along the x-axis; (6) model turning full 360° along the y-axis; and (7) model transitioning to front view. The hDyn1 helix is sky blue and actin filaments are in other colors. Scale bar, 10 nm.

**Supplementary Movie 11. Cryo-electron tomogram and modelling of hDyn1-actin interaction.**

The movie includes: (1) z-sections through the cryo-electron tomogram; and (2) modelling of actin filaments and hDyn1 helix on the background of the tomogram.

**Supplementary Movie 12. Cryo-electron tomogram of Shi and actin.**

Z-sections of cryo-electron tomogram generated from a tilt series collected on cryo grids containing Shi and actin filaments. Scale bar, 10 nm.

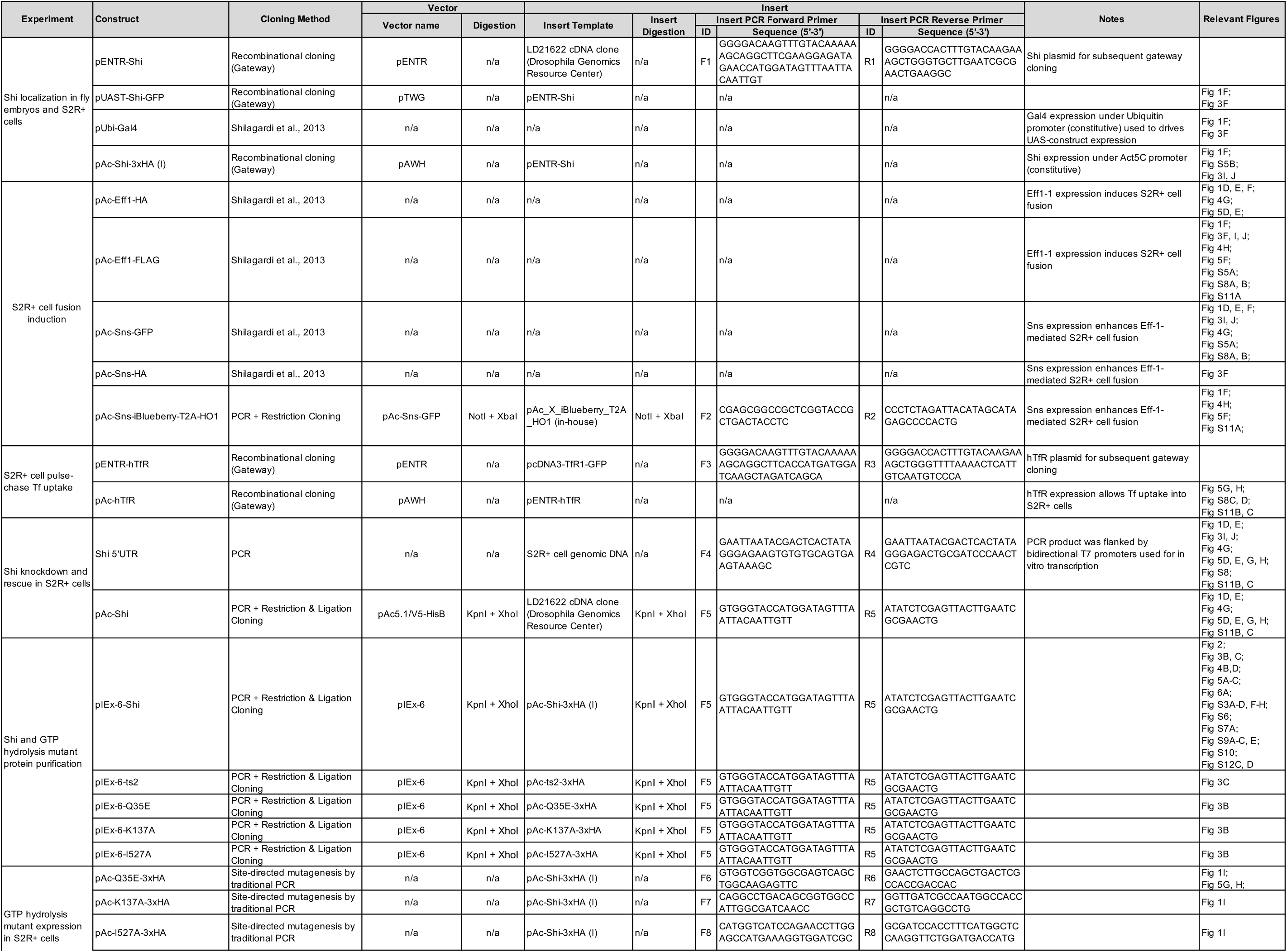
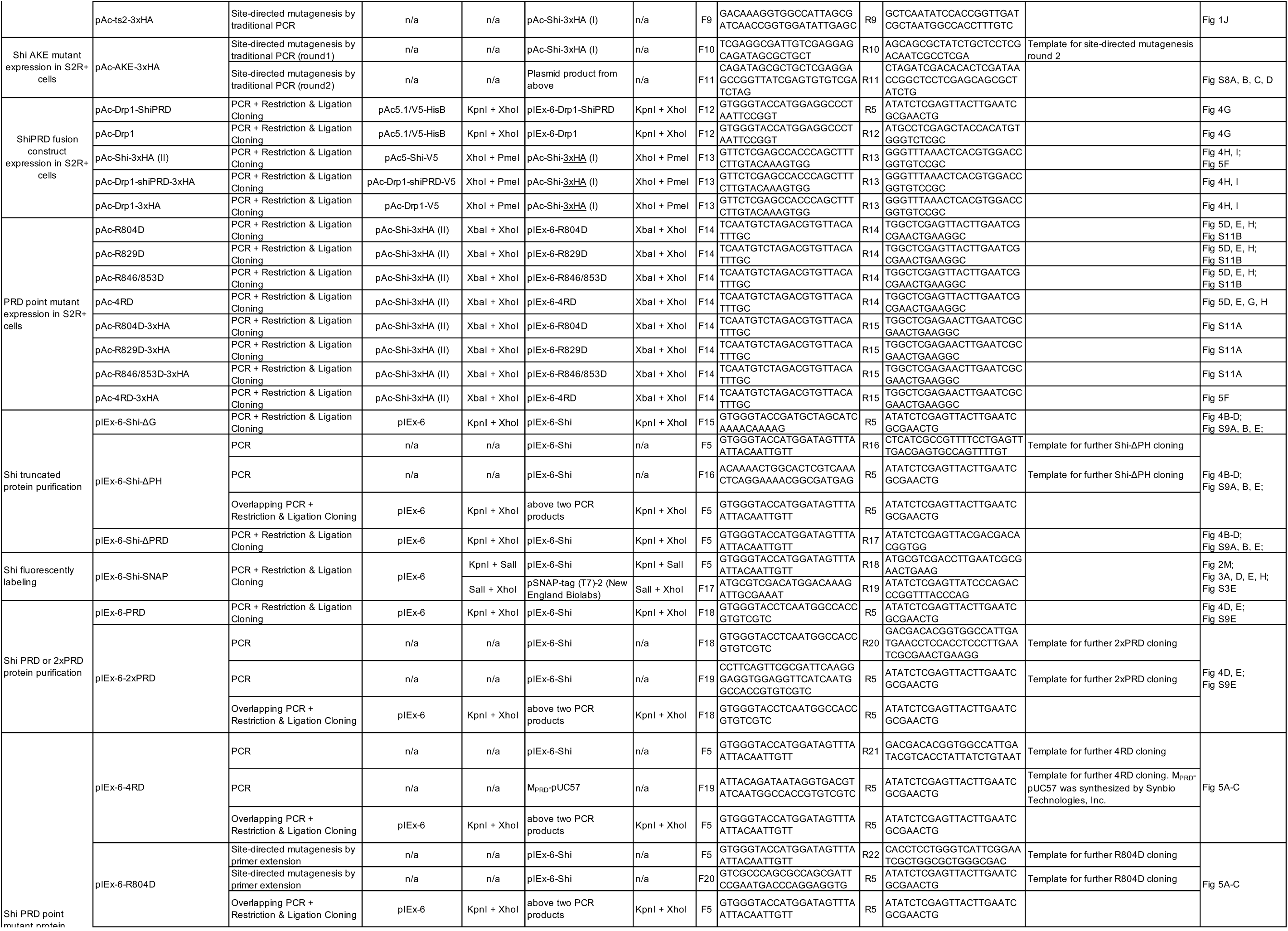
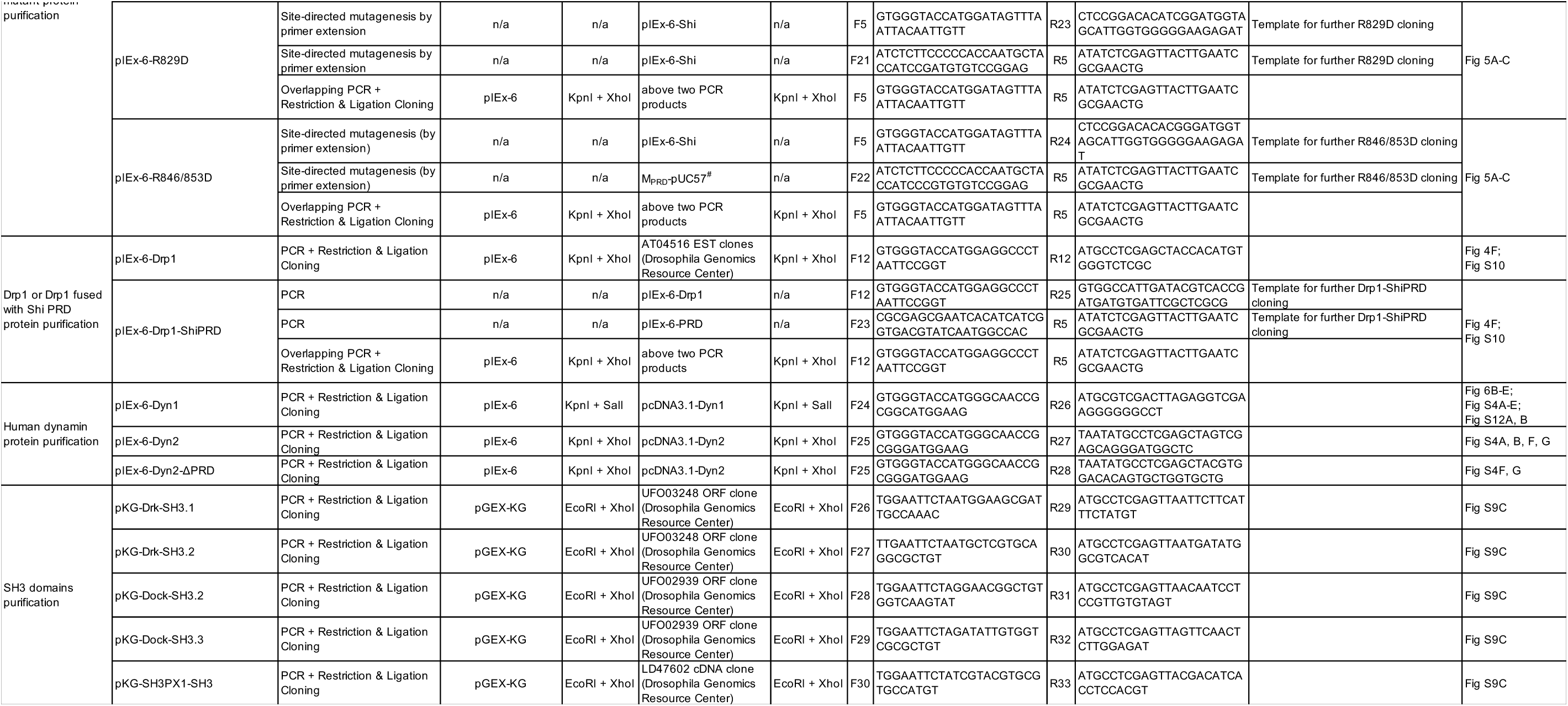

## Materials and Methods

### Fly stocks and genetics

The following stocks were obtained from the Bloomington *Drosophila* Stock Center: *w*^1118^ (wild type; BL#3605), *shi^ts1^* (BL#7068), *shi^ts2^* (BL#2248), *kette*^J4-48^ (BL#8753), *twi-GAL4* (BL#914), *UAS-Actin5CmRFP* (BL#24778), *UAS-Clc-GFP* (BL#7107), and *UAS*-*AP-2α-GFP* (*AP-2α*^MI06502-GFSTF.1^) (BL#59834). Other stocks used were: *sns^40-49^/CyO*^48^, *wasp^3^/TM3*^49^, *sltr^S1946^/CyO*^50^, *rP298-GAL4*^51^, and *sns-GAL4*^52^. Transgenic flies carrying *UAS-Shi-GFP* were generated by P-element-mediated germline transformation. To express Shi-GFP in all muscle cells, flies carrying the *UAS-Shi-GFP* transgene were crossed with *twi-GAL4*. For transgenic rescue experiments, *shi^ts2^; UAS-shi-GFP* flies were crossed with *shi^ts2^; twi-GAL4* (all muscle cells), or *shi^ts2^, rP298-GAL4* (founder cells), or *shi^ts2^; sns-GAL4* (FCMs) flies. For phenotypic analyses of embryos carrying *shi^ts1^* or *shi^ts2^* mutations, embryos were collected for 1 or 2 hrs at 20 °C and incubated for 16 or 15 hours at 18 °C prior to temperature shift. The embryos were then shifted to the restrictive temperature of 32 (or 34) °C for either 2 hours (or 4 hours), followed by fixation and immunostaining experiments to assess the F-actin foci phenotype (or myoblast fusion phenotype). Washes and dechorionation prior to fixation were carried out with buffers pre-warmed to 32 °C.

### Cell culture

S2R+ cells were cultured in Schneider’s medium (Gibco) supplemented with 10% fetal bovine serum FBS (Gibco) and penicillin/streptomycin (Sigma). S2R+ cells were induced to fuse as previously described^8^. Briefly, 1.2×10^6^ cells were plated into each well of a 6-well plate and transfected with Sns and Eff-1 constructs (200ng each) using Effectene (Qiagen) per the manufacturer’s instructions. Transfected cells were then fixed after 48 or 72 hrs for phenotypic analyses. For Shi RNAi knockdown (KD) experiments, cells were incubated with 3 µg/ml of dsRNA for four days, transfected with appropriate DNA constructs, and continued to be incubated with 3 µg/ml of dsRNA for 48-72 hrs. For rescue experiments, a Shi construct (100 ng) was co-transfected with Sns and Eff-1 in the Shi KD cells. For drug treatment, cells were incubated with 2 μM, 3 μM or 4 μM of MiTMAB (Abcam, ab120466) at 24 hrs post-transfection, and incubated for another 24 hrs before fixation.

Sf9 cells were cultured in suspension medium of Sf-900 III SFM (Gibco) with L-Glutamine (Gibco) and Antibiotic-Antimyotic (Gibco). The cells were grown at a density of 0.5 × 10^6^ − 6 × 10^6^ cells/ml with 150 rpm shaker at 27 °C. For protein expression, cells were diluted to 0.5 × 10^6^ cells/ml with fresh medium and cultured for 24 hrs, followed by transfection with corresponding DNA constructs using Transfection Reagent I (Avanti) per manufacturer’s instructions and continued culturing for 48 hrs.

HeLa cells (ATCC CCL-2) were cultured in DMEM (Gibco) supplemented with 1X non-essential amino acids (Gibco) and 10% FBS (Gibco).

### Immunofluorescence staining and confocal microscopy

For *Drosophila* embryo staining, embryos were dechorionated, rinsed, and fixed in 1:1 mix of heptane and 4% formaldehyde in PBS for 20 min, devitellinized in 1:1 heptane/methanol by vigorous shaking, washed in methanol, and incubated in PBSBT (0.2% BSA and 0.1%Triton in PBS) for 2 hrs, unless otherwise noted. Samples were incubated with primary antibodies overnight at 4 °C, followed by PBSBT washes and incubation with secondary antibodies. The following primary antibodies were used: rabbit anti-Shi (1:500)^53^, rabbit anti-MHC (1:2000; gift from Bruce Paterson), rat anti-Tropomyosin (1:1000; Abcam, ab50567), guinea pig anti-Duf (1:500)^7^, rabbit anti-Ants (1:2000)^54^; rabbit anti-Eve (1:30; Developmental Studies Hybridoma Bank (DSHB), 3C10), rabbit anti-GFP (1:500; Invitrogen, A11122), chicken anti-GFP (1:5000; Invitrogen, A10262). The following secondary antibodies were used at 1:200: Alexa488-, 568-, and 647-conjugated (Invitrogen) and biotinylated antibodies (Vector Laboratories) made in goat. Vectastain ABC kit (Vector Laboratories, PK-4000) and the TSA system (Perkin Elmer, SAT701001EA) were used to amplify weak fluorescent signals for anti-Eve staining. Nuclei were labeled with Hoechst 33342 (1:1000; Invitrogen, H3570). For Shi antibody staining, embryos were fixed in 1:1 mix of heptane and 37% formaldehyde for 5 min. For F-actin labeling, embryos were fixed in formaldehyde-saturated heptane for 1 hour at room temperature, hand-devitellinized, and incubated with Alexa488-, Alexa568- or Alexa647-conjugated phalloidin (1mg/ml) (Sigma) at 1:200 for 2 hrs at room temperature.

For S2R+ cell staining, cultured cells were fixed in 3.7% formaldehyde for 10 min, unless otherwise noted. Samples were incubated with primary antibodies for 1 hr at room temperature, followed by PBSBT washes and incubation with secondary antibodies for 1 hr at room temperature. The following primary antibodies were used: mouse anti-hTfR (1:600)^55^, rabbit anti-HA (1:600; Sigma, H6908), and hDyn2 (1:50; Santa Cruz Biotechnology, G4). Nuclei were labeled with Hoechst 33342 (1:1000; Abcam, 27671). The following secondary antibodies were used at 1:200: Alexa488-, 568-, and 647-conjugated (Invitrogen) and biotinylated antibodies (Vector Laboratories) made in goat. F-actin was labeled with Alexa488-, Alexa568- or Alexa647-conjugated phalloidin (Sigma) at 1:900.

Labeled samples were mounted in Vectashield (Vector Laboratories) or Aqua-Poly/Mount (Polysciences). Confocal microscopy was performed using a Zeiss LSM 700 microscope equipped with EC Plan Neofluar DIC M27 40x/1.3 and Plan Apochromat DIC 63x/1.4 oil objectives, a Nikon A1R confocal microscope (Ti-E; Nikon) equipped with CFI Plan Fluor 40x/1.3 and CFI Plan Apo VC 60x/1.4 oil objectives, and Leica TCS SPE confocal microscope (DMi8; Leica) equipped with an HC PL APO 20x/0.8 dry objective. The images were processed using Adobe Photoshop CS6 and ImageJ (NIH) softwares.

### Molecular biology

All molecular reagents generated and used in this study are described in Table S1.

### Electron microscopy

The high-pressure freezing and freeze substitution (HPF/FS) method was used to fix fly embryos as described^7,56^. Briefly, fly embryos at stage 12-14 were frozen with a Bal-Tec device, followed by freeze-substitution with 1% osmium tetroxide, 0.1% uranyl acetate in 98% acetone and 2% methanol on dry ice. Embryos were then embedded in Epon (Sigma), cut in thin sections (70 nm) with an ultramicrotome (Leica EM UC7), mounted on copper grids, and post-stained with 2% uranyl acetate for 10 min and Sato’s lead solution^57^ for 1 min for contrast enhancement. Images were acquired on a transmission electron microscope (CM120; Philips).

### Proteins purification

Most recombinant proteins generated in this study (*Drosophila* Shi, ts2, Q35E, K137A, I527A, Shi-SNAP, Shi-ΔG, Shi-ΔPRD, Shi-ΔPH, PRD, 2×PRD, Drp1, Drp1-ShiPRD, Shi-4RD, R804D, R829D, and R846/853D; human Dyn1, Dyn2 and Dyn2-ΔPRD) were expressed in inset Sf9 cells transiently transfected with the corresponding constructs unless otherwise noted. These N-terminal His-tagged fusion proteins were purified with nickel-nitrilotriacetic acid resin (Thermo) according to the manufacturer’s instructions. PRD and 2×PRD were further purified with gel filtration. All purified proteins were dialyzed against 20 mM HEPES, pH 7.3, 150 mM KCl, 1 mM EGTA and 1 mM DTT, flash frozen in liquid nitrogen, and stored at −80°C.

Non-tagged hDyn1 used in this study (Fig. S4A-E) was purified using GST-Amphiphysin II-SH3 domain as described ^58^. Non-tagged hDyn1 was used to compare with N-terminal His-tagged hDyn1 in the activity of actin bundling and no differences were found revealed by negative stain EM assay. Human Drp1 was purified using bacterial BL21 DE3 RIPL cells as described previously^59^.

All the SH3 domains used in this study (*Drosophila* Drk-SH3.1, Drk-SH3.2, Dock-SH3.2, Dock-SH3.3, SH3PX1-SH3, and amphiphysin II-SH3) were expressed in *Escherichia coli* BL21 (DE3). GST-tagged fusion proteins were affinity purified using Glutathione superflow agarose (Thermo) according to the manufacturer’s instructions. The GST-tags were removed by thrombin. All purified proteins were dialyzed against 10 mM Tris-HCl, pH 8.0, flash frozen in liquid nitrogen, and stored at −80°C.

### Co-sedimentation assay

High-speed and low-speed F-actin co-sedimentation assays were performed to determine the F-actin binding and bundling activity of Shi (and its mutants), respectively. 3 μM preassembled F-actin was incubated with the indicated concentrations of Shi or a mutant in 5 mM HEPES, pH 7.3, 50 mM KCl, 2 mM MgCl_2_, 0.5 mM EGTA, 3 mM imidazole, 60 nM ATP, 60 nM CaCl_2_, 0.4 mM DTT and 0.003% NaN_3_ for 30 min at room temperature. The samples were then spun for 30 min at 4 °C at 50,000g for high-speed and 13,600g for low-speed co-sedimentation, respectively. The supernatants and pellets resulting from centrifugation were resolved by 10% SDS-PAGE and stained with InstantBlue (expedeon).

### Total internal reflection fluorescence microscopy

To visualize Shi-actin bundles, 2 μM rabbit skeletal muscle actin (90% unlabeled and 10% rhodomine-actin) (Cytoskeleton Inc.), 1 μM Shi were incubated in 6.5 mM HEPES, pH 7.3, 50 mM KCl, 1 mM MgCl_2_, 0.3 mM EGTA, 80 nM ATP, 80 nM CaCl_2_, 0.3 mM DTT and 0.004% NaN_3_ for 2 hrs. 10 μl of the reaction mix was mixed with 2× TIRF buffer (1x TIRF buffer: 50 mM KCl, 1 mM MgCl2, 1 mM EGTA and 10 mM imidazole, 100 mM DTT, 0.2 mM ATP, 15 mM glucose, 20 μg/ml catalase, 100 μg/ml glucose oxidase, 0.2% BSA, 0.5% methylcellulose, pH 7.0), and loaded onto flow chambers prepared as follows. Flow chambers were coated with 10 nM NEM-myosin in HS-TBS buffer (50 mM Tris-HCl, pH 7.5 and 600 mM NaCl) for 1 min, washed with HS-BSA (1% BSA, 50 mM Tris-HCl, pH 7.5 and 600 mM NaCl) twice, washed with LS-BSA (1% BSA, 50 mM Tris-HCl, pH 7.5 and 150 mM NaCl) twice and incubated for 5 min, and washed with TIRF buffer. The loaded samples on the flow chambers were incubated for 5 min and subjected to TIRF imaging. Nikon Eclipse Ti microscopy equipped with an Apochromat DIC N2 TIRF 100x/ 1.49 oil objective lens and an Andor DU-897 X-8654 camera. Convolutedness is calculated as the ratio of traced filament length to the length of the longest side of a bounding rectangle encompassing the same filament^60^.

To visualize branched actin polymerization mediated by Arp2/3 on dynamin-actin bundles, both Shi-actin and human Dyn1-actin bundles were examined. Shi-actin bundles were preassembled with 2 μM Shi, 1 μM G-actin and 1 μM Alexa 488-conjugated phalloidin. The preassembled samples were loaded onto flow chambers, incubated for 5 min, washed and incubated with TIRF buffer with or without 1 mM GTP for an additional 5 min. Subsequently, 50 nM Arp2/3 (Cytoskeleton Inc.), 1 μM VCA (Cytoskeleton Inc.) and 250 nM rhodamine-actin were loaded onto flow chambers with or without 1mM GTP and subjected to TIRF imaging immediately. To visualize Shi in the actin polymerization process, Shi-SNAP was labeled fluorescently with SNAP-Surface 488 (New England Biolabs), and Shi-SNAP-Surface 488-mediated actin bundles were visualized by TIRF imaging. hDyn1-actin bundles were preassembled with 2 μM hDyn1, 1 μM G-actin and 1 μM Alexa 488-conjugated phalloidin. The preassembled samples were loaded onto flow chambers, incubated for 5 min, washed and incubated with TIRF buffer with or without 0.01 mM GTP for an additional 1 min. After another brief wash with TIRF buffer, 50 nM Arp2/3, 1 μM VCA and 250 nM rhodamine-actin were loaded into flow chambers without GTP and subjected to TIRF imaging immediately.

To monitor the process of Shi bundling of preassembled actin filaments, experiments were performed on His-tagged Ezrin-bound supported lipid bilayers (SLBs). SLBs were prepared as previously described ^61^. Briefly, 96-well glass-bottomed plate (Matrical) was pretreated with Hellmanex III (Hëlma Analytics) overnight, thoroughly rinsed with MilliQ H2O, washed with 300 μl of 6 M NaOH for 30 min at 45°C and repeated once, rinsed with 500 μl of MilliQ H2O for three times, and washed with 500 μl of HEPES buffer (50 mM HEPES, pH 7.3, 150 mM NaCl and 1 mM TCEP) for three times to equilibrate wells and finally 200 μl of HEPES buffer was kept in each well. 10 μl of SUVs (98% POPC, 2% DOGS-NTA and 0.1% PEG-5000PE) were added into wells and incubated for 1 hr at 40 °C to form SLBs, followed by three washes with 500 μl of HEPES buffer. Next, 100 μl of BSA buffer (50 mM HEPES, pH 7.3, 50 mM KCl, 1 mM TCEP, 1 mM MgCl_2_ and 1 mg/ml BSA) was added into each well and incubated for 30 min at room temperature. Ultimately, His-tagged Ezrin at 40 nM in BSA buffer was added into each well and incubated for 1 hr at room temperature, followed by five washes with 500 μl of BSA buffer to remove unbound His-Ezrin. Preassembled actin filaments (90% unlabeled actin and 10% rhodamine-actin) was added into wells and incubated for 5 min at room temperature followed by 100 μl of image buffer addition (0.02 mg/ml catalase, 0.1 mg/ml glucose oxidase and 15 mM glucose in BSA buffer) to adjust the actin concentration to 100 nM. Immediately before imaging, Shi was added into a well to a final concentration of 10 nM and imaged by a TIRF/iLAS2 TIRF/FRAP module (Biovision) mounted on a Leica DMI6000 microscope base equipped with a 100x/1.49 oil objective lens and a Hamamatsu ImagEMX2 EM-CCD camera.

To monitor Shi-mediated actin bundling in the process of actin polymerization, experiments were performed in flow chambers prepared as previously described ^62^ except that piranha cleaning was replaced with UVO treatment. Briefly, microscope slides and coverslips (#1.5; Fisher Scientific) were washed for 30 min with acetone and for 10 min with 95% ethanol, sonicated for 2 hrs with Helmanex III detergent (Hellma Analytics), washed extensively with milliQ water and treated with UVO cleaner (Jetlight Company) for 10 min. Slides were then immediately incubated for 18 hrs with 1 mg/ml mPeg-Silane (5,000 MW) in 95% ethanol (pH 2.0). Parallel strips of double-sided tape were placed on a coverslip to create multiple flow chambers. To initiate the experiment, 1.5 μM Mg-ATP-actin (10% tetramethylrhodamine (TMR)-labeled) and 1 μM Shi was mixed with 2X TIRF buffer [1X TIRF buffer: 10mM imidazole (pH 7.0), 50 mM KCl, 1 mM MgCl2, 1 mM EGTA, 50 mM DTT, 0.2 mM ATP, 50 μM CaCl2, 15 mM glucose, 20 μg/ml catalase, 100 μg/ml glucose oxidase, and 0.2% (400 centipoise) methylcellulose] and transferred to a flow chamber for imaging. TIRF microscopy images were collected at 10-sec intervals using a Nikon Inverted Ti-E microscope base equipped with a 100x/1.49 oil objective lens and an Evolve EMCCD camera (Photometrics) with through-the-objective TIRF illumination.

### Negative stain electron microscopy

Negative stain electron microscopy (EM) was performed to visualize the distribution of dynamin and mutants along actin bundles. 1 μM actin were incubated with the indicated concentrations of appropriate proteins in 6.5 mM HEPES, pH 7.3, 50 mM KCl, 2 mM MgCl_2_, 0.3 mM EGTA, 20 nM ATP, 20 nM CaCl_2_, 0.3 mM DTT and 0.001% NaN_3_ at 4°C overnight. The reaction mixture was loaded onto carbon-coated, glow discharged 400 mesh copper grids and incubated for 5 min, followed by two quick washes in actin polymerization buffer (50 mM KCl, 1 mM MgCl_2_, 1 mM EGTA and 10 mM imidazole). The grid was stained in two drops of 0.75% uranyl formate for 20 sec and air-dried. Electron micrographs were collected on a JEOL 1200-EX II transmission electron microscope.

To test the effects of GTP on dynamin-actin bundles by negative stain EM, 1 μM actin were incubated with the indicated concentrations of dynamin or mutants in the presence or absence of 1 mM GTP at room temperature for 30 min.

To visualize dynamin helices on lipid nanotubes, lipid tubes were generated with 40% phosphatidylcholine (1,2-dioleoyl-sn-glycero-3-phosphocholine, DOPC), 40% NFA-GalCer, 10% cholesterol and 10% PtdIns(4,5)P2 (to a total lipid concentration of 1 mM) (Avanti). 3 μM dynamin or a mutant was incubated with lipid nanotubes in lipid buffer (20 mM HEPES, pH 7.3, 150 mM KCl, 2 mM MgCl_2_ and 1 mM DTT) at 4°C overnight, followed by negative stain described above except that the wash buffer was done in the lipid buffer.

To visualize dynamin self-assembled rings, 0.2 mg/ml dynamin was dialyzed into 20 mM HEPES, pH 7.3, 25 mM KCl, 1 mM EGTA and 1 mM DTT at 4°C overnight. Samples were loaded onto grids and incubated for 1 min, followed by two quick washes with the same dialysis buffer and staining in two drops of 0.75% uranyl formate for 1 min.

### Super resolution microscopy

To visualize the distribution of Shi along actin bundles generated by purified proteins, Shi-SNAP-Surface 488 was incubated with actin at different molar ratios ([Shi]: [actin] = 1:8, 1:1 or 2:1) and an equimolar concentration of Alexa568-conjugated phalloidin as actin with or without 1 mM GTP, in 6.5 mM HEPES, pH 7.3, 50 mM KCl, 1 mM MgCl_2_, 0.3 mM EGTA and 0.3 mM DTT for 2 hrs at room temperature. Subsequently, the samples were diluted 20-fold in fluorescence buffer (0.5% methycellulose, 50 mM KCl, 1 mM MgCl2, 1 mM EGTA, 10 mM imidazole, 100 mM DTT, 0.02 mg/ml catalase, 0.1 mg/ml glucose oxidase and 15 mM glucose, 0.1 mM CaCl2, 0.1 mM ATP and 0.005% NaN3). 4 μl of diluted samples were mounted onto a slide, covered with poly-L-lysine coated coverslip, and subjected to SIM imaging.

To visualize Shi localization along actin bundles at the fusogenic synapse, 1.2×10^6^ S2R+ cells were seeded onto each well of a 6-well plate and transfected using Effectene (Qiagen) with either pAc-Eff1-HA (150 ng), pAc-Sns-iBlueberry-T2A-HO1 (150 ng) pUAST-Shi-GFP (50 ng) and pUbi-Gal4 (50 ng) for SIM imaging, or pAc-Eff1-FLAG (150 ng), pAc-Sns-HA (150 ng), pUAST-Shi-GFP (50 ng) and pUbi-Gal4 (50 ng) for STED imaging. Samples were washed twice with PBS supplemented with 1 mM CaCl_2_ and 0.5 mM MgCl_2_ (PBS++) at 36-48 hrs post transfection, then fixed with 4% paraformaldehyde diluted in cytoskeleton preservation buffer (80 mM PIPES pH6.8, 5mM EGTA, 2mM MgCl_2_) for 10 min at 37°C^63^. Samples were washed in PBS, permeabilized and incubated in PBSBT, followed by immunostaining. For SIM imaging, the following labeling reagents were used: rabbit anti-HA antibody (1:600; ThermoFisher, 715500), Alexa488- or Alexa647-conjugated goat anti-rabbit antibody (1:600; Invitrogen) and Alexa568-conjugated phalloidin (1:200; Sigma). For STED imaging, the following labeling reagents were used: rabbit anti-HA antibody (1:600; ThermoFisher, 715500), mouse anti-GFP antibody (1:600; DSHB, 8H11), CF405M-conjugated goat anti-rabbit antibody (1:600; Biotium), Star580-conjugated goat anti-mouse antibody (1:600; Abberior) and StarRed-conjugated phalloidin (1:200; Abberior).

To visualize hDyn2 localization along actin bundles on comet tails, 4.5×10^5^ HeLa cells were seeded onto 25 mm glass coverslips (Warner Instruments, Cat 64-0715), deposited in 6-well plates the day before infection. GFP-expressing *Listeria monocytogenes* strain 10403s (provided by Dr. Dan Portnoy, UC Berkeley) was cultured in 3 ml of brain heart infusion (BHI) media (Gibco) at 30°C overnight without shaking until OD_600_ = 0.8 was reached. The following day, 1 ml *Listeria* culture was centrifuged at 16,000xg for 1 min, washed with PBS, and resuspended in a final volume of 1 ml PBS. HeLa cells were infected with *Listeria* at a multiplicity of infection (MOI) of 60, and infection synchronized by centrifugation for 10 min at 800g. The cells were then incubated at 37°C for 90 min, washed with PBS, followed by another incubation in fresh media containing 50 µg/ml gentamicin for 5 hours. HeLa cells were washed twice with PBS supplemented with 1 mM CaCl_2_ and 0.5 mM MgCl_2_ (PBS++), then fixed with 4% paraformaldehyde diluted in cytoskeleton preservation buffer (80 mM PIPES pH6.8, 5mM EGTA, 2mM MgCl_2_) for 10 min at 37°C^63^. Samples were washed in PBS, permeabilized and incubated in PBSBT, followed by immunostaining. The following labeling reagents were used: mouse anti-hDyn2 (1:50; Santa Cruz Biotechnology, G4), Star580-conjugated goat anti-mouse antibody (1:600; Abberior) and StarRed-conjugated phalloidin (1:200; Abberior).

SIM imaging was performed using a Nikon N-SIM E microscope (Ti-E; Nikon), equipped with a CFI SR Apochromat TIRF 100x/1.49 oil objective lens and an ORCA-Flash 4.0 sCMOS camera (Hamamatsu Photonics K.K.). Excitation of the fluorophores was performed with three lasers (488/561/640nm). STED imaging was performed using an Abberior Instruments Expert Line STED microscope (Abberior Instruments, Göttingen, Germany), equipped with an SLM based easy3D module, an Olympus IX83 microscope body and an UPLSAPO 100x/1.4 oil objective. Excitation of the fluorophores was performed with pulsed excitation lasers at 561 nm and 640 nm. A pulsed 775 nm laser (pulse width, ∼1.2 ns) was used for STED. Both lasers were synchronized and operated at 40 MHz. Time-gated detection of the fluorescence was performed at 605-625nm for the orange channel and 650-720 nm for the deep red channel. The images were processed using Adobe Photoshop CS6 and ImageJ (NIH) software.

### Fluorescence recovery after photobleaching

To visualize the dynamics of Shi along actin bundles with or without GTP, actin bundles were prepared by incubating 1 μM Shi-SNAP-Surface 488 and 1 μM actin in 6.5 mM HEPES, pH 7.3, 50 mM KCl, 1 mM MgCl_2_, 0.3 mM EGTA and 0.3 mM DTT for 2 hrs at room temperature. Before imaging, 0.25 μM Shi-SNAP-Surface 488 in fluorescence buffer with or without 1 mM GTP were mixed with the samples. To perform FRAP assay, two pre-photobleaching images were recorded, followed by photobleaching of the regions of interest (ROI) and quickly bleached to ∼15% of its original intensity. Subsequently, images were acquired every 30 sec by a Nikon A1R confocal microscope (Ti-E; Nikon) equipped with a CFI SR Apochromat TIRF 100x/1.49 oil objective lens. The mean fluorescence intensity of the photobleached region was measured by ImageJ software and unbleached region was used to correct the curve.

### GTPase activity assay

The GTPase activity of Shi was measured according to previously published methods^64^. For basal GTPase assay, 1 μM Shi was incubated in GTPase assay buffer (GAB-150; 20 mM HEPES, pH 7.5, 150 mM KCl, 2 mM MgCl_2_ and 1 mM DTT) with a series of GTP concentrations (25, 50, 100, 200, 400, 800 and 1200 μM) at room temperature. Samples were harvested from the reaction mixes at 5-min interval over a 30 min time period, and reactions were stopped by EDTA addition to a final concentration of 100 mM. Malachite green solution was added to each sample and the absorbance at 650 nm was determined using a microplate reader. Released Pi detected by malachite green was fitted to a linear curve vs. time course, and the GTP hydrolysis rate was calculated for each GTP concentration. Subsequently, the GTP hydrolysis rates were plotted against the initial concentrations of GTP to calculate the Michaelis-Menten constants K_cat_ and K_m_.

For actin-stimulated GTPase assay, 0.1 μM Shi was pre-incubated with 0.3 μM actin in GTPase assay buffer-50 (GAB-50; 20 mM HEPES, pH 7.5, 50 mM KCl, 2 mM MgCl_2_ and 1 mM DTT) for 1 hr to form stable Shi-actin bundles at room temperature. GTP was added to reaction mixes to reach a series of final concentrations of 10, 15, 25, 50, 100, 150 and 200 μM. Samples were harvested from the reaction mixes at 2-min interval over a 12 min time period, and processed as described above.

### Western blot

Expression constructs were transfected in S2R+ cells. Cells were harvested, washed in PBS and lysed in lysis buffer (50mM Tris pH 7.5, 200 mM NaCl, 10 mM MgCl2, 1% NP-40, 0.5% Triton X-100) supplemented with 1x protease inhibitor cocktail (Sigma). After centrifugation, cell lysates were analyzed on an SDS-PAGE gel. The following antibodies were used in blotting: rabbit anti-HA (1:1000; ThermoFisher, 715500), mouse anti-tubulin (1:1000; DSHB, E7), chicken anti-GFP (1:5000; ThermoFisher; A10262), anti-rabbit-HRP (1;10,000; Invitrogen), anti-mouse-HRP (1;10,000; Invitrogen), and anti-chicken-HRP (1:20,000; Invitrogen).

### Pulse-chase transferrin uptake assay

1.2×10^6^ S2R+ cells were plated in each well of a 6-well plate, transfected with 100 ng of human TfR (and 100ng of rescue construct if applicable) using the Effectene (QIAGEN) according to manufacturer’s instructions. At 36 hrs post-transfection, cells were washed with PBS, incubated in PBS++++ (1mM CaCl_2_, 1mM MgCl_2_, 5mM Glucose and 0.2% BSA) for 30 min, and incubated with 5 µg/ml Alexa488- or Alexa568-conjugated transferrin (Tf) (Molecular Probes, T13342 and T23365) for 5 min. Subsequently, cells were placed on ice, washed three times with cold PBS, three times with acetic acid (0.2M acetic acid, 0.2M NaCl at pH 2.5) for 5 min each, three times again with cold PBS, and fixed with 4% formaldehyde, and stained for hTfR and HA. Stained cells were mounted and subjected to confocal imaging.

For image analysis, a custom-written ImageJ (Fiji) macro was used. In brief, for each image analyzed, transfected cells were identified and manually outlined, yielding the total surface area (TSA). Then, within these TSAs, fluorescent signals of interest were automatically determined using the triangular threshold method, and particles were analyzed though the build-in ImageJ algorithm. Relative Tf uptake was determined by Tf particle count divided by the TSA, followed by normalization to the relevant control within the dataset.

### Cryo-electron tomography

Cryo-ET was used to determine the 3-dimensional organization of actin filaments bundled by dynamin. 5 μM actin was incubated with 0.625 μM of either hDyn1 or Shi in 20 mM HEPES, pH 7.3, 50 mM KCl, 1 mM EGTA, 80 nM ATP, 80 nM CaCl_2_, 1 mM DTT and 0.004% NaN_3_ at 4°C overnight. Aliquots of 3.5 μl of the overnight actin/dynamin reaction mixture with 10 nm gold fiducials was loaded onto glow-discharged gold C-flat grids (Electron Microscopy Sciences, CF-1.2/1.-4Au-50) for 20 sec, back-blotted on the sample-free side with filter paper for 2 sec (22 °C, 90% humidity) and then plunged into liquid ethane using the EM Grid Plunger (Leica Microsystems). Electron micrographs for the cryo-ET study were collected with a 200 kV TF20 transmission electron microscope (FEI) at 29,000x magnification, with a defocus of −3.0 μm using a K2 summit camera (Gatan). Tilt series was collected using SeriaEM at angles between ∼−60° and +60° for each data set. Tomograms were generated from the tilt series with the IMOD package^65^. Modelling of dynamin (EMDB code: EMDB-7957) and actin (provided by Dr. Krishna Chinthalapudi using the PDB entry 5ONV) into tomograms was done in Chimera^66^, and was also guided by segmented tomograms obtained using the EMAN2 semi-automated segmentation package^67^. The helical parameters of dynamin were calculated based on the diameter, pitch and known distance between dynamin subunits along the helical path.

## Acknowledgements

We thank the Bloomington stock center for fly stocks, Dr. Bruce Paterson for the MHC antibody, Dr. Melissa Mikolaj for help with generating the model in iMOD, Drs. Haifeng He and Huaibin Wang for help with generating the cryo-tomograms using SerialEM, and Dr. Kishna Chinthalapudi for providing the actin filament for modeling in Chimera. This work was supported by NIH grants (R01 AR053173 and R01 GM098816), American Heart Association Established Investigator Award, and HHMI Faculty Scholar Award to E.H.C.; NIH grant (R01 AI083359), HHMI and Simons Foundation, and Welch Foundation (I-1704) to N.M.A.; NIH grant (R01 GM127673), Chan Zuckerberg Biohub Investigator Award, and HHMI Faculty Scholar Award to A. F.; NIH grant (R01 GM42455) and Welch Foundation (I-1823) to S.L.S. M.K.R is an HHMI investigator. R. Z. was supported by an American Heart Association postdoctoral fellowship; N. G. by an American Heart Association predoctoral fellowship; D.M.L. by a Canadian Institute of Health Research postdoctoral fellowship; and J.A.D by a National Research Service Award from NIDDK (F32 DK101188).

## Author contributions

S.K. initiated the project. R.Z., N.G., D.M.L., S.K and E.H.C. design the project, performed experiments and discussed the data. R.Z. carried out biochemistry, negative stain EM, and in vitro imaging experiments in Fig. 2 (except for 2E, F), Fig. 3 (except for 3F, G), Fig. 4 (except for 4G, H), Fig. 5A-C, Fig. 6A, Fig. S3, 4, 6, 7, 9 and 10. N.G. carried out fly genetics and cell culture experiments in Fig. 1A, B, G and H, Fig. 5G and H, Fig. S1, Fig. S2F, Fig. S8. D.M.L carried out cell culture and super resolution imaging experiments in Fig. 1D-F, Fig. 3F and G, Fig. 4G and H, Fig. 5D-F, Fig. S5, Fig. S11. S.K performed live imaging and immunostaining experiments in Fig. 1C and Fig. S2A-E. R.Z. and E.H.C. collaborated with J.J. and J.H on the cryo-ET experiments in Fig. 6 and Fig. S12, with J.W. and M.G. on TIRF imaging experiment in Fig. 2E and F, with R.K. and A.F. on experiments with human Drp1 in Fig. S10, and with J.D. and M.R. on live imaging of actin bundling in Fig. 2D. N.G., D.M.L. and E.H.C. collaborated with M.M. and S.S. on the endocytosis experiments in Fig. 5G, Fig. S8D and Fig. S11B. D.M.L. and E.H.C. collaborated with M.A. and N.A. on the actin comet tail experiments in Fig. 3G and Fig. S5B. D.L. and E.H.C. collaborated with G.Z. on the EM experiments in Fig. 1I. P.K. worked with D.M.L. in generating constructs used in Fig. 1F, Fig. 5D-F and Fig. S11B. P.P. worked with R.Z. on the negative stain EM experiments in Fig. 2–6, Fig. S3, S4 and S10. B.R. worked with R. Z. on the lipid tubule experiments in Fig. 2J, Fig. S4E and Fig. S9B. J.K. worked with N.G. on foci invasion measurement in Fig. 1G. R.Z., N.G., D.M.L. and E.H.C. made the figures. R.Z., D.M.L. and E.H.C. wrote the paper. All authors commented on the manuscript.

## Data availability

The main data supporting the findings of this study are available within the article and its Supplementary Information files. All other data supporting the findings of this study are available from the corresponding author upon reasonable request.

## Author information

The authors declare no competing financial interests.

